# Cristae formation is a mechanical buckling event controlled by the inner membrane lipidome

**DOI:** 10.1101/2023.03.13.532310

**Authors:** Kailash Venkatraman, Christopher T. Lee, Guadalupe C. Garcia, Arijit Mahapatra, Daniel Milshteyn, Guy Perkins, Keun-Young Kim, H. Amalia Pasolli, Sebastien Phan, Jennifer Lippincott-Schwartz, Mark H. Ellisman, Padmini Rangamani, Itay Budin

## Abstract

Cristae are high curvature structures in the inner mitochondrial membrane (IMM) that are crucial for ATP production. While cristae-shaping proteins have been defined, analogous mechanisms for lipids have yet to be elucidated. Here we combine experimental lipidome dissection with multi-scale modeling to investigate how lipid interactions dictate IMM morphology and ATP generation. When modulating phospholipid (PL) saturation in engineered yeast strains, we observed a surprisingly abrupt breakpoint in IMM topology driven by a continuous loss of ATP synthase organization at cristae ridges. We found that cardiolipin (CL) specifically buffers the IMM against curvature loss, an effect that is independent of ATP synthase dimerization. To explain this interaction, we developed a continuum model for cristae tubule formation that integrates both lipid and protein-mediated curvatures. The model highlighted a snapthrough instability, which drives IMM collapse upon small changes in membrane properties. We also showed that CL is essential in low oxygen conditions that promote PL saturation. These results demonstrate that the mechanical function of CL is dependent on the surrounding lipid and protein components of the IMM.

**Synopsis:**

- critical lipidic breakpoint for yeast mitochondria phenocopies the loss of cristae-shaping proteins in the IMM.
- saturation controls membrane mechanical properties and modulates ATP synthase oligomerization.
- mitochondrial-specific lipid cardiolipin can functionally compensate for increased phospholipid saturation and is required for cristae formation in low oxygen environments.
- mathematical model for cristae membrane tubules predicts a snapthrough instability mediated by both protein and lipid-encoded curvatures.

**Synopsis Figure:** 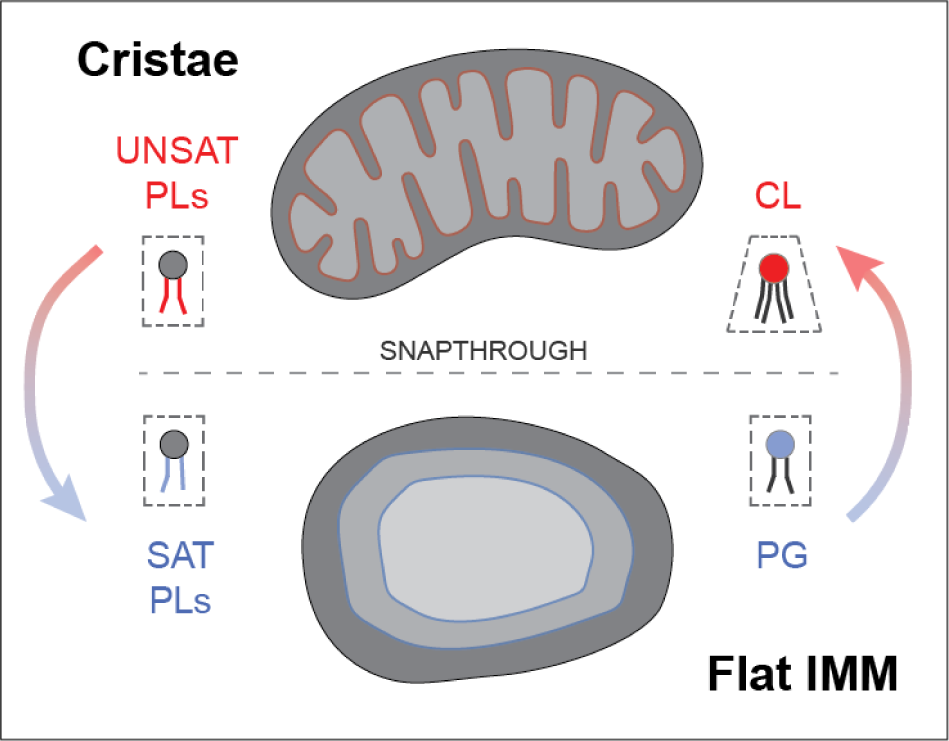

## Introduction

Mitochondria are ubiquitous eukaryotic organelles whose membrane architecture is required for their metabolic and non-metabolic functions (Nunnari & Suomalainen, 2012). The inner mitochondrial membrane (IMM) is the site of the Electron Transport Chain (ETC) and consists of two regions defined by their curvature: the flat inner boundary membrane (IBM), adjacent to the outer mitochondrial membrane (OMM), and cristae membranes (CM), which invaginate into the matrix and are connected to the IBM by crista junctions (CJs) (Daems & Wisse, 1966; Perkins *et al*, 1997). CM structure is dependent on organism, tissue and physiological state (Perkins *et al*, 1997; Revel *et al*, 1963) but is commonly composed of tubular and lamellar membranes (Revel *et al*, 1963; Mannella, 2006b; Zick *et al*, 2009; Pánek *et al*, 2020; Mendelsohn *et al*, 2022). Cristae effectively increase membrane surface area for ETC reactions and could also act as a ‘proton sink’ where protons travel expeditiously to F_1_F_0_ adenosine-triphosphate (ATP) synthase (Davies *et al*, 2011; Rieger *et al*, 2014; Cogliati *et al*, 2016). Cristae could also act as diffusion barriers for metabolites between the intracristal space (ICS) and the intermembrane space (IMS), controlling the flux of ADP/ATP through the adenine nucleotide translocase (ANT) (Mannella *et al*, 1994; Mannella *et al*, 1997; Frey & Mannella, 2000; Mannella, 2006a). All these potential functions of cristae are dependent on their high intrinsic membrane curvature.

The IMM is shaped by proteins that drive assembly and maintenance of cristae. The molecular determinants of CM are best understood in *Saccharomyces cerevisiae,* where ATP synthases form ribbon-like rows of dimers which induce curvature along tubule and lamellar rims (Dudkina *et al*, 2005; Strauss *et al*, 2008; Davies *et al*, 2011; Blum *et al*, 2019). Loss of the dimerization subunit *g*, Atp20p, results in monomeric ATP synthases and onion-like mitochondria with flat layers of IMM that run parallel with the OMM (Arnold *et al*, 1998; Paumard *et al*, 2002; Arselin *et al*, 2004; Rabl *et al*, 2009). Reconstituted ATP synthase dimers spontaneously assemble into rows driven by changes to elastic membrane bending energies (Anselmi *et al*, 2018) and are sufficient to form tubular liposomes (Blum *et al*, 2019). At the CJ, the dynamin-related GTPase optic atrophy protein 1 (OPA1)/Mgm1p interact with the mitochondrial contact site and cristae organizing system (MICOS) complex (Frezza *et al*, 2006; Harner *et al*, 2011; Hoppins *et al*, 2011; Patten *et al*, 2014; Glytsou *et al*, 2016; Hu *et al*, 2020). Cells lacking Mgm1p feature a completely flat IMM (Sesaki *et al*, 2003; Harner *et al*, 2016), while loss of the major MICOS subunit Mic60p results in elongated cristae sheets that do not contain CJs (Rabl *et al*, 2009; Harner *et al*, 2011).

In addition to their proteinaceous determinants, mitochondrial lipids are hypothesized to play key roles in shaping cristae. The predominant phospholipids (PLs) of the IMM are phosphatidylethanolamine (PE), phosphatidylcholine (PC) and cardiolipin (CL) (Zinser *et al*, 1991; Mejia & Hatch, 2016). IMM phospholipids, or their phosphatidylserine (PS) precursors in the case of PE, are imported from the ER at contact sites (Horvath & Daum, 2013), CL and its precursor phosphatidylglycerol (PG) are synthesized in and remain localized to the IMM. Among PLs, CL is unique in featuring four acyl chains whose larger cross sectional area contributes to an overall conical shape (LeCocq & Ballou, 1964; Beltrán-Heredia *et al*, 2019). In liposomes, CL localizes to regions of high curvature and can drive pH-dependent invaginations (Khalifat *et al*, 2008, 2011; Ikon & Ryan, 2017), suggesting a role in promoting curved membrane topologies. The curvature of CL itself varies depending on the local lipid and chemical environments (Chen *et al*, 2015; Beltrán-Heredia *et al*, 2019) molecular simulations predict key roles for its ionization state (Dahlberg & Maliniak, 2010) and binding of counter ions (Konar *et al*, 2023). Despite these biophysical data, the fundamental mitochondrial functions of CL are not fully resolved. In the genetic disorder Barth syndrome, loss of the acyl chain remodeler Taffazin causes reduced amounts and altered composition of CL (Adès *et al*, 1993; Bione *et al*, 1996) leading to abnormal cristae (Acehan *et al*, 2007), which have also been observed in cell lines lacking CL synthesis (Claypool & Koehler, 2012; Ren *et al*, 2014; Ikon & Ryan, 2017; Paradies *et al*, 2019). In yeast, however, loss of cardiolipin synthase (Crd1p) does not render a respiratory or morphological phenotype under regular growth temperatures (Jiang *et al*, 1997; Baile *et al*, 2014). It thus remains unknown if CL serves a mechanical role in the IMM, or has more organism-specific functions relating to ETC enzymes (Xu *et al*, 2021) and their organization (Zhang *et al*, 2005).

The acyl chain composition of mitochondrial PLs also differs from other organelles (Harayama & Riezman, 2019) and broadly regulates membrane biophysical properties. The IMM is enriched in unsaturated and polyunsaturated PLs, which promote membrane fluidity, and lacks sterols and saturated sphingolipids, which promote membrane ordering (Filippov *et al*, 2003; Vance, 2015). During synthesis, lipid unsaturation is controlled by the activity of fatty acid desaturases, such as Ole1p in yeast (Bard, 1972; Stukey *et al*, 1989). *OLE1* was discovered in genetic screens for both unsaturated fatty acid auxotrophy and mitochondrial distribution and morphology genes (MDM). Mutations in *mdm2* resulted in abnormal mitochondrial morphology and inheritance (McConnell *et al*, 1990; Stewart & Yaffe, 1991) but were later identified as *OLE1* alleles, indicating an unexplained link between desaturase activity and mitochondrial structure. In mammalian cells, addition of exogenous saturated fatty acids such as palmitic acid (PA) drives mitochondrial dysfunction (Sparagna *et al*, 2000; Penzo *et al*, 2002; Jheng *et al*, 2012) and can cause the progressive loss of CMs (Xue *et al*, 2019). Metabolic diseases, such as obesity and type 2 diabetes, have also been associated with both saturated fat accumulation and mitochondrial stress (Petersen *et al*, 2004; Lowell & Shulman, 2005).

Here we combine experimental perturbations with mutli-scale modeling to elucidate new roles for conserved mitochondrial lipids in IMM morphology, using yeast as a model system. We first used genetic manipulation of PL saturation and observed a surprising lipidic breakpoint, in which the IMM becomes flat and mitochondria lose their ATP synthesis capacity. This transition is controlled by the IMM lipidome in two distinct ways: through modulation of ATP synthase oligomerization by PL saturation and through loss of intrinsic membrane curvature provided by CL. We develop a mathematical model to explain these effects by considering the energetics of lipid and protein-mediated membrane curvature. We then show CL function is dependent on growth conditions that modulate PL saturation, most notably oxygenation, and that it has an essential role in natural yeast growth environments.

## Results

### Systematic modulation of the yeast PL double bond profile reveals a critical mitochondrial breakpoint in IMM curvature and ATP generation

Bulk PL membrane properties are in part controlled by the stoichiometry between saturated and unsaturated acyl chains. To modulate lipid saturation in budding yeast, we utilized a library of promoters controlling the expression of Ole1p (Figure 1A). We focused on four strains, saturated fatty acid (SFA) 1-4, which showed a range in lipid saturation (Figure 1A). SFA1 features a wild-type (WT) PL composition while SFA2-4 have consecutively increasing levels of PL acyl chain saturation due to lower levels of *OLE1* expression (Appendix Figure S1A). Among PC and PE lipids, WT and SFA1 strains possess predominantly di-unsaturated PLs, while SFA 2, 3 and 4 (weaker *OLE1* expression) show an increasing ratio of mono to di-unsaturated species and incorporation of fully saturated PLs (Figure 1A). We observed potentially compensatory adaptations to increasing saturation in whole cell PLs (Appendix Figure S1), including a decrease in the PE/PC ratio (Appendix Figure S1B), used by several organisms to increase membrane fluidity (Janssen *et al*, 2000; Dawaliby *et al*, 2016), an increase in PI (Appendix Figure S1B), and shortening of acyl chains length (Appendix Figure S1C), which also occurs during yeast cold adaptation (Al-Fageeh & Mark Smales, 2006).

**Figure 1:**
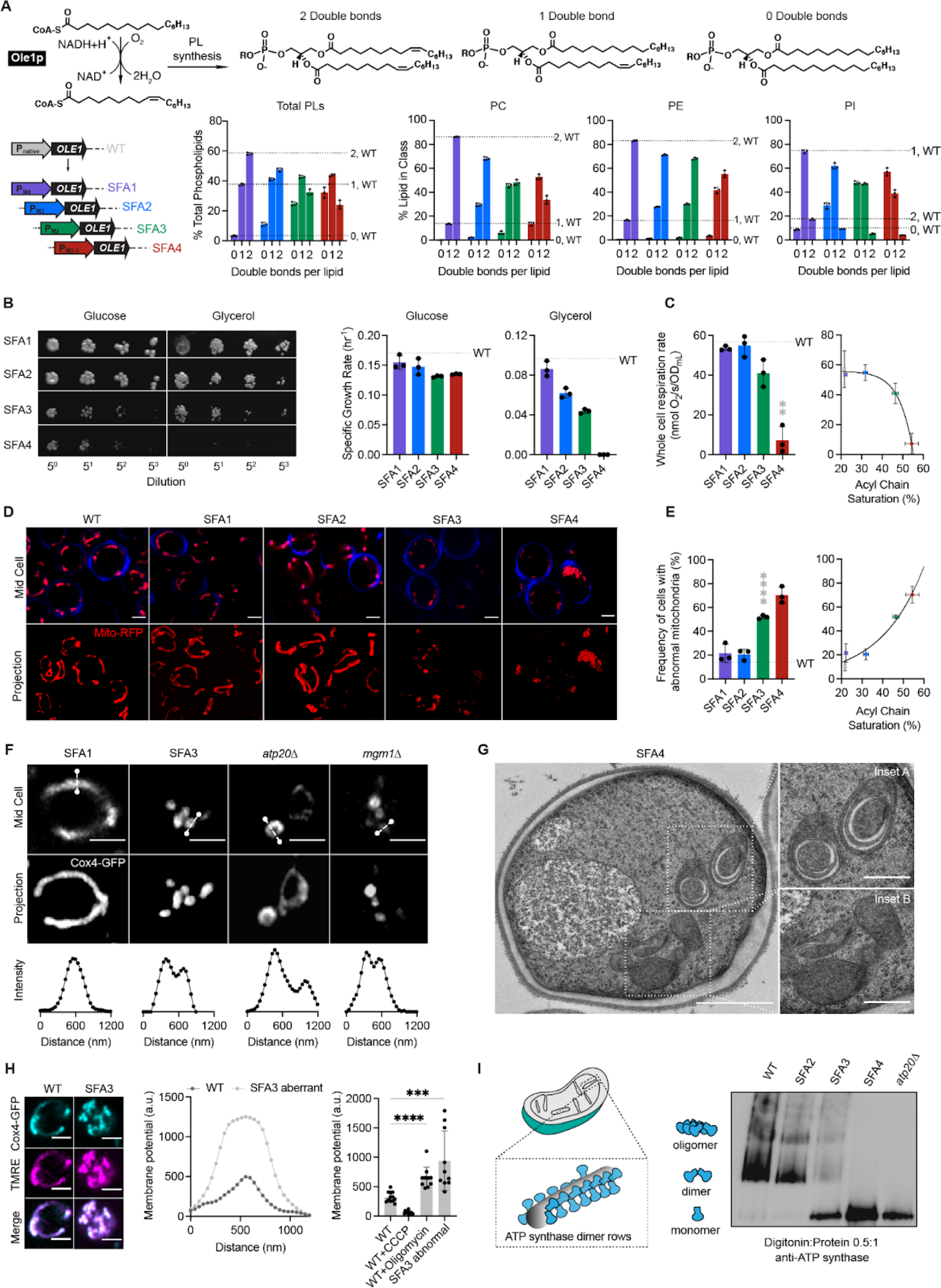
Modulation of *OLE1* expression results in a critical level of PL saturation driving loss of ATP synthase oligomerization and mitochondrial morphology. **(A)** The yeast desaturase, Ole1p, is an oxygen-dependent enzyme that introduces *cis* double bonds at the C9 position (top left). SFA strains were generated via promoter substitution, resulting in progressively decreasing levels of *OLE1* expression (bottom left). Lipidomics analysis showing double bond distributions of the total PL pool as well as of individual PLs within SFA strains; the wild-type distribution is depicted with dotted lines. Error bars indicate SD from biological replicates n=3. **(B)** SFA4 cells lose viability under respiratory conditions. Shown are serial dilutions of yeast cells plated on media containing fermentable (glucose) and non-fermentable (glycerol) carbon sources and specific growth rates for each in liquid cultures. Error bars indicate SD from n=3 independent cultures. **(C)** (Left) SFA3 and SFA4 cells show a drop in whole cell respiration, measured using a Clark electrode. Error bars indicate SD from n=3 independent cultures. **p=0.004, unpaired two-tailed t-test of SFA4 compared against wild-type. (Right) Scatter plot depicting a fitted single exponential (R^2^ =0.99) for the decrease in respiration as a function of acyl chain saturation. **(D)** SFA3 and SFA4 cells lose tubular mitochondrial morphology. Shown are representative Airyscan confocal micrographs of yeast expressing matrix-localized RFP (mts-RFP). Cells were stained with cell wall-binding calcofluor white (blue) for clarity. Scale bars, 2 μm. **(E)** Mitochondrial morphology changes between SFA2 and SFA3 strains, as assayed by confocal microscopy of cells harboring an mts-RFP plasmid. N>50 cells were counted in biological triplicate in each condition. Error bars indicate SD from n=3 independent cultures. ****p < 0.0001, unpaired two-tailed t-test of SFA3 compared against wild-type. (Right) Scatter plot depicting a fitted single exponential (R^2^ =0.97) for the increase in frequency of abnormal mitochondria as a function of acyl chain saturation. **(F)** Airyscan confocal micrographs of yeast expressing IMM protein Cox4-GFP showing hollow mitochondria in SFA3 cells, as is also observed in mutants of CM shaping proteins. Scale bars, 2 μm. Profiling analysis (below) depicts fluorescence intensity as a function of the distance across the indicated mitochondrion; two peaks indicate a lack of fenestrated IMM. **(G)** Thin section TEM micrographs of high-pressure frozen (HPF) SFA4 yeast showing the appearance of an onion-like (Inset A) IMM and total loss of CM (Inset B). Scale bars, 1 μm (full image), 400 nm (insets A and B). **(H)** Abnormal mitochondria show an increase in membrane potential, consistent with loss of ATP synthase, but not ETC activity. (Left) Representative micrographs are shown of individual cells expressing Cox4-GFP and stained with TMRE. (Center) Example membrane potential line scan plots showing an example of an abnormal SFA3 cell with higher TMRE intensity compared to WT. (Right) Quantification of TMRE peak intensities from line profiling analysis of N>10 cells per condition showing a higher membrane potential in aberrant mitochondria, as well as those where ATP synthase is inhibited by oligomycin (5 μM), and lower potential in cells treated with the uncoupler CCCP (20 μM). The line profile measured the intensity between two points crossing an individual mitochondria. Scale bars, 2 μm. ***p<0.0005, ****p<0.0001 unpaired two-tailed t-test of SFA3 aberrant and WT+oligomycin compared against WT. **(I)** ATP synthase oligomerization is lost with increasing PL saturation. SFA2 mitochondria lose higher order oligomers observed in WT cells, while SFA3 and SFA4 mitochondria possess predominantly monomeric ATP synthase, similar to *atp20*Δ cells. Shown are BN-PAGE western blots of digitonin-solubilized purified mitochondria, *atp20*Δ is the monomeric ATP synthase control.

When handling these strains, we observed that SFA3 and SFA4 cells were characterized by loss of viability in non-fermentable carbon sources (Figure 1B), oxygen consumption (Figure 1C), and tubular mitochondrial networks (Figure 1D-E, Appendix Figure S2). In contrast, non-mitochondrial physiology – assayed by growth under fermentation conditions (Figure 1B), morphology of other organelles (Figure EV1A), and activation of the unfolded protein response (UPR) (Figure EV1B) – remained unaffected within this range of *OLE1* mutants. Thus, a modest increase in saturation conferred a sudden loss of mitochondrial function. We further observed that SFA3 and SFA4 mitochondria featured gaps in Cox4-GFP, a subunit of cytochrome *c* oxidase in the IMM, similar to *atp20*Δ and *mgm1* cells that lack CMs (Figure 1F). Transmission electron microscopy (TEM) analysis of SFA4 cells prepared by high-pressure freeze substitution (HPFS) (Figure 1G, Appendix Figure S3A,B) showed mitochondria with flat IMM membranes similar to those in *atp20*Δ (Paumard *et al*, 2002) or *mgm1* cells (Harner *et al*, 2016). The moderate increase in saturation in SFA2 cells, which does not cause aberrancy independently, also showed epistasis with loss of cristae shaping proteins at both the cristae ridge (Atp20p) and rim (Mic60p of the MICOS complex) (Figure EV2A, EV2B). We thus hypothesized that PL saturation has a specific effect on formation of CMs.

We next asked how increasing lipid saturation resulted in the aberrant mitochondrial morphologies. SFA3 mitochondria retained normal levels of ETC complexes as well as respiratory chain supercomplexes (SCs) (Figure EV2C). Aberrant mitochondria in SFA3 cells retained membrane potential (Figure 1H) as measured by staining with tetramethylrhodamine ethyl ester (TMRE), which was also inconsistent with a loss of proton-pumping reactions in the ETC. Further examination showed that TMRE fluorescence in aberrant mitochondria was elevated to a similar level as WT cells treated with the ATP synthase inhibitor oligomycin, suggesting that they were characterized by an impediment to ATP synthesis itself. We analyzed ATP synthase in isolated mitochondria with blue native PAGE (BN-PAGE) western blotting, which can assay the organization of complexes. In mitochondria from SFA strains, increasing saturation progressively altered ATP synthase organization. WT mitochondria contained predominantly dimeric and oligomeric ATP synthase, while oligomers are lost in SFA2 cells. SFA3 and SFA4 mitochondria predominantly contained monomeric ATP synthases, similar to *atp20*Δ cells (Figure 1I). SFA3 cells notably retained other cristae-shaping proteins, including Mic60p and the long and short forms of Mgm1p (Figure EV2D), indicating that this effect on ATP synthase was specific. Furthermore, all phenotypes observed in SFA3 mitochondria, including an increase in the number of mtDNA nucleoids (Figure EV2E) and the average contact distance with the ER (Figure EV2F), could also be phenocopied by loss of ATP synthase dimerization in *atp20*Δ. These data suggested that saturated lipids directly modulate ATP synthase organization and through this mechanism alter CM formation.

### Modeling of transport processes in aberrant mitochondria suggest a mechanism for loss of ATP generation

Loss of ATP synthase oligomerization provides a mechanism for loss of CM morphology, but did not fully explain the respiratory phenotypes of SFA3/4 and *atp20*Δ because monomeric ATP synthases retain ATPase activity (Arnold *et al*, 1998; Paumard *et al*, 2002). To explore the functional consequences of CM loss, we more extensively characterized aberrant IMM morphologies (Figure 2). Two types of structures were observed in wide-field TEM micrographs of SFA4 cells: onion-like mitochondria with multiple IMM layers, which were observed in at least 40% of cells, and flat IMMs with a single membrane running parallel to the OMM (Appendix Figure S3A). Tomography of onion-like IMMs showed connection vertices and alternating layers of matrix and IMS (Appendix Figure S3B), as previously observed in *atp20*Δ cells (Paumard *et al*, 2002), suggesting a continuous IMM that could arise during membrane growth (Appendix Figure S3C) to resemble discrete IMM layers. We carried out 3D reconstructions of these morphologies from multi-tilt TEM (Appendix Figure S4A) using the GAMer 2 platform (Lee *et al*, 2020) and computed two types of local membrane curvature across their surfaces, based on the maximum principal curvature (*k*1) and minimum curvature (*k*2) at each point. The mean between these two principal curvatures (*k*1 and *k*2) provides information about how the surface normal changes at a given point (mean curvature), while the difference between them captures the extent of anisotropy between the two directions (deviatoric curvature). This analysis showed that the IMM of SFA2 mitochondria were marked by regions of high (>25 μm^-1^) mean and deviatoric curvatures (Figure 2A, Figure 2C-D), but these were completely absent in both types of aberrant morphologies (Figure 2B, Figure 2C-D).

**Figure 2:**
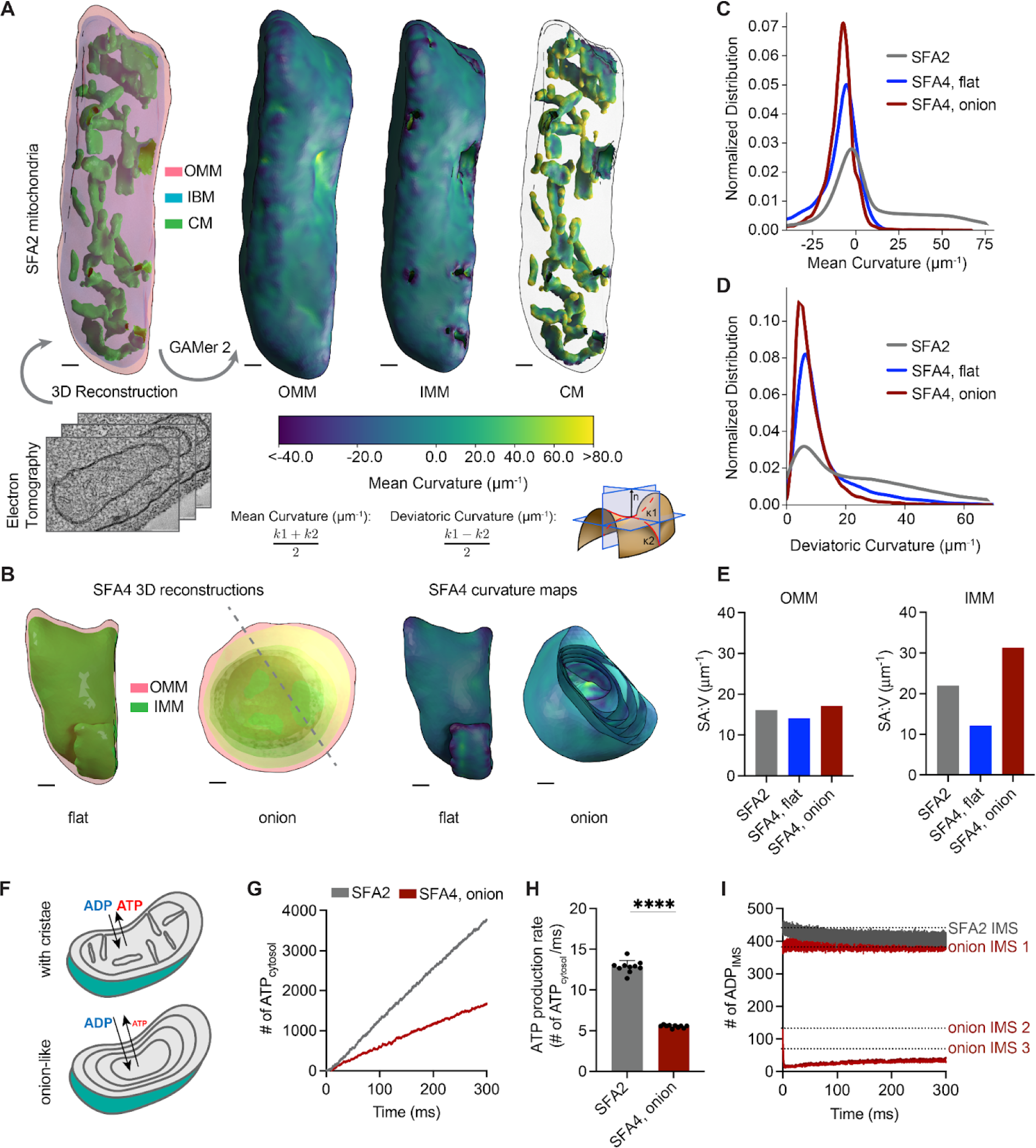
PL saturation shifts the IMM to a regime of low membrane curvature, which reduces IMM surface area and modeled transport rates needed for ATP generation. **(A)** 3D reconstructions of mitochondrial membrane topology and quantification of curvature using the GAMer 2 pipeline. Shown are electron tomograms of mitochondria from SFA2 cells, which show regular, tubular CM. CMs are highlighted alongside the OMM and the inner boundary membrane (IBM) in the 3D reconstruction. (Right) Maps of mean curvature of OMM, IMM and CM computed using GAMer2. Scale bars, 50 nm. **(B)** 3D reconstructions of SFA4 mitochondria showing flat (*mgm1*Δ*-*like) and onion (*atp20*Δ*-*like) abnormal morphologies. Also shown are the maps of mean curvature showing the IMM of these abnormal mitochondria, using the same color scale as in A. The onion-like IMM is sliced at an angle to illustrate the many layers; each layer is spherical in nature. Scale bars, 50 nm. **(C)** Histograms of mean curvature distributions of inner membranes generated from SFA2 and SFA4 reconstructions, highlighting that high mean curvature areas in SFA2 IMM are lost in SFA4 cells. **(D)** Histograms of deviatoric curvature distributions of inner membranes generated from SFA2 and SFA4 reconstructions, which are mechanical corollaries to curvature induced by ATP synthase dimerization, highlight that high deviatoric curvature areas in SFA2 IMM are lost in SFA4 cells. **(E)** The IMMs of onion mitochondria show increased surface area:volume (SA:V) ratio compared to SFA2 while flat mitochondria show a decrease in the SA:V without major changes to the OMM. **(F)** Schematic of ATP production in mitochondria containing normal CMs and those showing a multi-layer, online-like IMM, highlighting how multiple membrane layers could impede in trafficking of ADP and ATP. **(G)** Modeled cytosolic ATP generation from mitochondria with a CM-containing morphology (taken from SFA2 tomogram) vs. an onion-like (taken from SFA4 tomogram) IMM. ATP generated in the cytosol in each condition is an average from 10 simulations. Details of the model equations and simulations are provided in the Appendix. **(H)** Comparison of cytosolic ATP generation rates derived from multiple Monte Carlo simulations shown in G. 10 simulations were run for each morphology. Error bars indicate SD; ****p<0.0001, unpaired two-tailed t-test. **(I)** Modeled substrate depletion in onion-like mitochondria. In the CM-containing mitochondrion, the ADP level in the IMS remains constant at ∼440 ADP throughout the 300 ms simulation. In the onion-like SFA4 mitochondrion, ADP remains constant only in the first IMS layer and is rapidly depleted in layers 2 and 3, indicating that the multi-layer structure could be limiting for ATP/ADP trafficking. Dotted lines represent initial values of ADP in each IMS layer.

Flat and onion-like IMMs differed dramatically in their surface area, with the former showing a reduced surface area:volume ratio (SA:V) and the latter an increased one. Previous modeling studies have shown that SA:V is an important determinant of flux of molecules between different compartments (Rangamani *et al*, 2013; Cugno *et al*, 2019; Calizo *et al*, 2020). We hypothesized that the online-like IMM structure would impede transport of ATP out of, and ADP into. To explore this hypothesis, we employed an MCell-based reaction-diffusion simulation pipeline for modeling ATP generation using EM-derived morphologies (Garcia *et al*, 2019). These simulations showed that mitochondria with multiple IMM layers cause lower ATP generation when cytosolic ADP concentration was kept constant (Figure 2F, Figure 2G-H). We also observed that predicted ATP production was inhibited by depletion of ADP as a substrate in the inner layers of the onion-like IMM, highlighting that ATP/ADP exchange by ANT could be limiting for high surface area IMM morphologies with low membrane curvature (Figure 2I). These simulations suggest that efficient substrate transport could be dependent on CMs and explain why low curvature IMM morphologies that retain high surface area could still impede ATP generation.

### Lipid saturation modulates membrane mechanical properties which are buffered by mitochondrial-specific headgroup changes

We next sought to define the lipid-encoded membrane properties that drive CM loss. We first analyzed the lipidomes from mitochondria isolated from SFA strains by density ultracentrifugation (Appendix Figure S5). For our analyses of the IMM we used whole mitochondrial lipidomes, as these were nearly identical to those of isolated IMMs from the same sample, with the notable exception of CL levels (Appendix Figure S1F). As in the whole cell lipidome, decreasing Ole1p activity in SFA strains increased mitochondrial PL saturation, but fewer fully saturated species were observed. Instead, the lipidic breakpoint in SFA3 cells corresponded to changes in the double bond distribution from predominantly di-unsaturated to monounsaturated PLs, e.g. PE-16:1/18:1 to PE-16:0/16:1 (Figure 3A-B), a surprisingly modest shift given the magnitude of the morphological and functional change.

**Figure 3:**
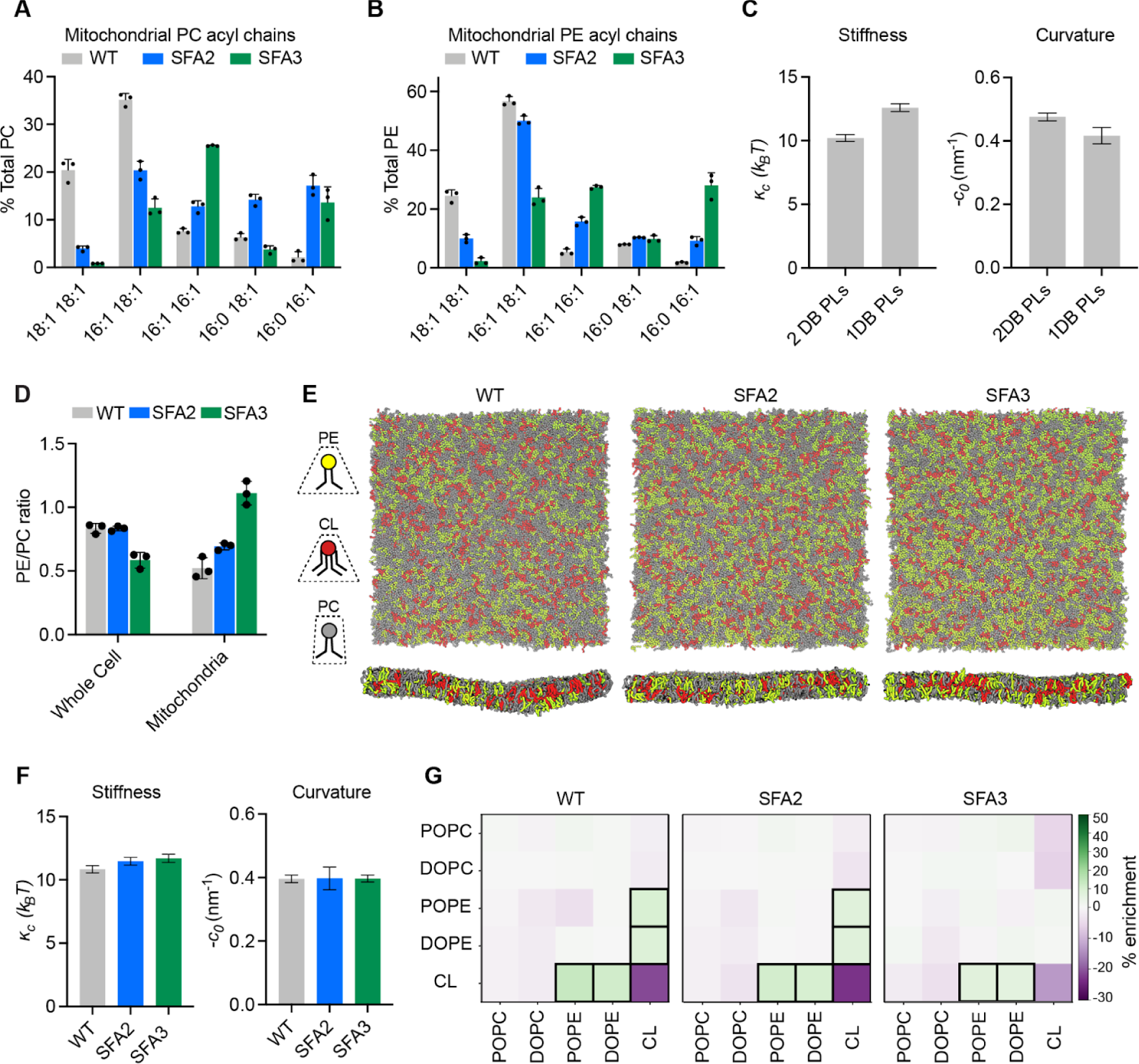
Modeling of lipid-driven mechanical changes in the IMM and their homeostatic responses. **(A)** Lipidomic profile of isolated mitochondria showing a transition from di-unsaturated to monounsaturated PC, n=3 biological replicates. Error bars represent SD. **(B)** Mitochondrial PE shows a similar transition from di-unsaturated species to monounsaturated species, n=3 biological replicates. Error bars represent SD. **(C)** Changes to membrane mechanical properties predicted by Martini 2 CG-MD simulations of mitochondrial-like lipid mixtures that shift from di-unsaturated (2 double bonds, DB) to mono-unsaturated (1 DB) PLs. Stiffness (bending modulus) increases while negative spontaneous curvature, derived from the first moment of the lateral pressure profile and bending modulus, decreases. Compositions of these ‘ideal’ systems are shown in Figure EV3A. **(D)** Increasing lipid saturation reduces the PE/PC ratio in whole cells, but increases it in isolated mitochondria, suggesting a curvature-based adaptation to increasing saturation. **(E)** Top-down and side-on snapshots of CG-MD bilayers showing headgroup adaptation to increasing saturation in SFA strains. **(F)** Simulations of the lipid systems derived from SFA mitochondria that incorporate homeostatic head-group changes. In contrast to the systems in (C), the complex systems show only small changes to membrane mechanical properties, suggesting that headgroup adaptations may offset the changes to mechanical properties associated with increased saturation. **(G)** Compositional enrichment within the local neighborhood of individual lipid types. Given a lipid type (y-axis) the color indicates the percent enrichment in likelihood of observing an x-axis labeled lipid within a 1.5 nm radius neighborhood compared to random distribution. Boxes indicate conditions with greater than 5% enrichment from random. PO/DOPE are enriched around CL, indicating the association between conically shaped lipids.

To explore the effects of this change in acyl chains on membrane properties, we employed coarse-grained molecular dynamics (CG-MD) simulations of lipid bilayers. CG-MD forgo atomistic detail in favor of improved sampling of complex mixtures (Marrink & Tieleman, 2013; Marrink *et al*, 2019), so we first tested whether they capture known mechanical properties of saturated PLs. For example, the stiffness (*к_c_*) of saturated PL bilayers is higher than those with unsaturated (Appendix Table S1A-B) and polyunsaturated chains (Rawicz *et al*, 2000; Manni *et al*, 2018). A second key parameter is the monolayer spontaneous curvature, *c_0_*, a measure of lipid shape. Cylindrical lipids, like PC, feature *c_0_* values near zero, while conical lipids like PE or CL feature negative *c_0_* values (Dymond, 2021). PL acyl composition also modulates *c_0_*, with more voluminous unsaturated chains favoring negative spontaneous curvatures (Szule *et al*, 2002). We simulated large membranes (∼40 nm x 40 nm, 5000 lipids) containing a simplified mixture of IMM lipids (50:30:20 PC:PE:CL) with either monounsaturated or diunsaturated PC & PE. Thermally induced height undulations were analyzed to derive *к*_c_ from each membrane composition. In parallel, small membranes (∼15 nm x 15 nm, 700 lipids) of identical compositions were used to compute the lateral pressure profiles, whose first moment is equal to the product of *к*_c_ and *c_0_*. Using *к_c_*values derived from the large systems, *c_0_* values can thus be extracted. The shift from diunsaturated to monounsaturated PLs led to a ∼30% increase in stiffness, *к*_c_, (Figure 3C), consistent with previous experimental measurements in monocomponent liposomes (Appendix Table S1A-B). Similarly, monounsaturated mixtures showed a ∼25% increase in *c*_0_, consistent with previous measurements by small angle x-ray scattering (Appendix Table S1C). We thus concluded that CG-MD using the Martini 2 force field can reproduce the expected changes to mechanical properties modulated by PL lipid saturation.

We then proceeded to analyze complex mixtures derived from mitochondrial lipidomes. In SFA strains, we observed a mitochondrial-specific change in headgroup composition: PE levels increased with increasing saturation at the expense of PC, resulting in an increase in the PE/PC ratio (Figure 3D, Appendix Figure S1E). In contrast, the PE/PC ratio decreased in the corresponding whole cell lipidome (Figure 3D, Appendix Figure S1B). We also observed an increase in PS levels, which serve as an intermediate for PE synthesis by Psd1p (Voelker, 1997). Because PE has a higher melting temperature than PC (Dawaliby *et al*, 2016), its increase argues against a fluidity-specific stress of saturation on the mitochondria. Instead, the high negative spontaneous curvature of PE suggested an adaptation to membrane spontaneous curvature itself. To understand the biophysical basis for this adaptation, we employed the CG-MD workflow described on lipid mixtures that mimic the changes in head-group composition in SFA strains (Figure 3E). In these systems, shifting from the WT mitochondrial lipidomes to those of the SFA3 mitochondria resulted in only a modest increase in stiffness (*к*_c_) and no change to the magnitude of *c*_0_, despite the increase in PL saturation (Figure 3F). These findings suggested that the mitochondrial specific increase in conical lipids act to buffer membrane mechanical properties relevant to curvature generation.

Beyond changes in the PE/PC ratio, our membrane simulations highlighted the key role of CL in dictating properties relevant to curvature generation. Replacement of CL with its precursor lipid PG resulted in an overall stiffening of simulated IMMs and loss of spontaneous curvature (Figure EV3D-E). Switching CL from the dianion form, predominant at neutral pH, to the monoanion form had the opposite effect, dramatically softening the bilayer and increasing spontaneous curvature (Figure EV3C-E). In simulations without CL, the increase in PE in SFA strains could only partially compensate for curvature lost when CL was removed (Figure EV3F). CL preferentially clustered with PE lipids in the simulated bilayers (Figure 3G), highlighting how conical lipids sort together in areas of high curvature (Callan-Jones *et al*, 2011). This association was maintained in SFA lipidomes, though the extent of CL-PE association was reduced in the saturated SFA3 bilayers (Figure 3G).

### An epistasis between PL saturation and CL abundance underlies IMM structure

The observation that mitochondria in SFA2 cells responded to increased PL saturation by increasing the abundance of high-curvature PE and CL lipids compared to WT (Appendix Figure S1D) led us to consider the interplay of PL curvature and saturation. While PL saturation itself can modulate intrinsic lipid curvature (*c_0_*) (Appendix Table S1C), there is no evidence that acyl chain composition differs across the two leaflets of the IMM, which would be needed to generate net membrane curvature across the bilayer (*C_0_*). We instead hypothesized that the effects of CL on predicted membrane curvature and the established asymmetric localization of CL in the IMM (Gallet *et al*, 1997) could provide membrane curvature independent of cristae shaping proteins, such as ATP synthase dimers. To explore this possibility, we tested out compositions based on previously measured asymmetries of CL in the yeast IMM. The simulations predicted that enrichment of CL in the IMM outer leaflet increased *c*_0_ and reduced *к*_c_ (Figure EV3G). Based on values for the former, we estimated that CL asymmetry could contribute at least a −0.05 nm^-1^ net bilayer curvature to the IMM (*C_0_*) (Figure EV3G).

We next asked if the IMM curvature imparted by CL could compensate for curvature lost by saturated PLs and loss of ATP synthase oligomerization. To test this hypothesis, we evaluated how loss of cardiolipin synthase, Crd1p, affects mitochondrial morphology in SFA1-4 strains (Figure 4A). Lipidomics confirmed that CL was absent in *crd1*Δ strains and PL saturation was unaltered (Figure 4A). As previously observed (Baile *et al*, 2014), *crd1*Δ cells did not show defects in mitochondrial morphology or cellular respiration in WT (*CRD1*) or SFA1 backgrounds (Figure EV4A-B). However, loss of CL had a dramatic phenotype in the SFA2 background, which has increased PL saturation and reduced ATP synthase oligomerization but normal mitochondrial morphology (Figure 4B-E). SFA2*crd1*Δ exhibited low oxygen consumption rates (Figure 4B), aberrant mitochondria (Figure 4C), and loss of CMs (Figure 4D-E), similar to SFA3/SFA4. Loss of morphology and respiration in SFA2*crd1*Δ was rescued by supplementation with oleic acid (OA), demonstrating that *crd1*Δ has a specific interaction with PL saturation (Figure EV4C).

**Figure 4:**
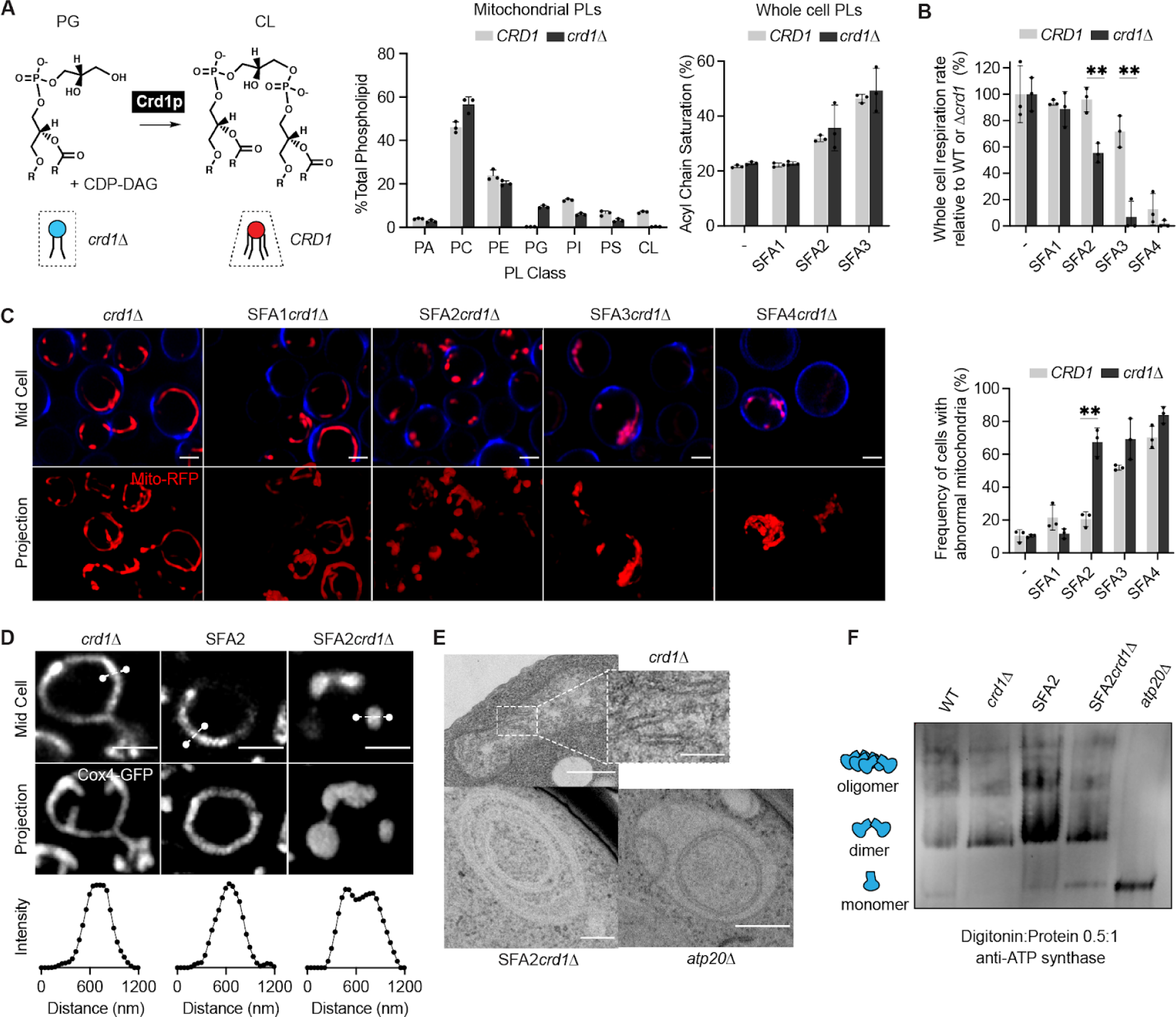
Epistasis between PL saturation and CL synthesis in shaping mitochondrial morphology. **(A)** Loss of Crd1p decreases mitochondrial CL content, but otherwise does not significantly affect the mitochondrial lipidome. (Left) Reaction schematic depicting cylindrical PG and CDP-DAG converted into conical CL by Crd1p. (Right) Head-group stoichiometry of lipidomes from isolated mitochondria from WT and *crd1*Δ cells and acyl chain saturation (% of acyl chains with a double bond) for *CRD1* and *crd1*Δ cells across the SFA series. Error bars indicate SD from n=3 biological replicates. **(B)** Loss of CL causes a loss of respiration in SFA2 cells. Whole cell respiration rates are shown for SFA strains, comparing *CRD1* with *crd1*Δ cells. Respirometry was conducted in biological triplicates (n=3) by Clark electrode. Error bars indicate SD. **p<0.005 unpaired two-tailed t-test between SFA2 and SFA3 strains and their corresponding *crd1*Δ mutants. **(C)** The morphological breakpoint shifts to SFA2 upon loss of CL. (Left) Representative Airyscan confocal micrographs of yeast expressing matrix-localized RFP (mts-RFP). Cells were stained cell wall-binding calcofluor white (blue) for clarity. Scale bars, 2 μm. (Right) Frequency of mitochondrial abnormality, N>50 cells were counted in n=3 independent cultures for each condition. Error bars indicate SD. **p=0.0011 unpaired two-tailed t-test between *CRD1* and *crd1* Δ in the SFA2 background. **(D)** SFA2*crd1* cells show hollow mitochondria, imaged with Cox4-GFP. Scale bars, 2 μm. Line profile analysis (below) depicts fluorescent intensity across the indicated mitochondria. **(E)** Thin section TEM micrographs of *crd1*Δ show IMM with cristae invaginations (see inset, scale bars 100 nm), while SFA2*crd1*Δ cells show an onion-like IMM similar to what has been observed in *atp20*Δ. Scale bars, 400 nm. **(F)** SFA2*crd1*Δ retains ATP synthase dimerization. Shown are BN-PAGE western blots run from digitonin-solubilized isolated mitochondria of SFA strains and WT with and without CL, *atp20*Δ is the monomeric ATP synthase control.

While loss of CL modulated CM formation under increased PL saturation, it did so independently of ATP synthase dimerization: *crd1*Δ cells showed no defect in ATP synthase oligomerization compared to *CRD1* cells, while SFA2*crd1*Δ cells maintained an identical level of oligomerization found in SFA2 cells (Figure 4F). However, loss of CL still showed a strong epistasis with the saturation induced loss of ATP synthase oligomerization in SFA2 cells, as well as with complete loss of dimerization in *atp20*Δ cells (Figure EV4D). Thus, CL acts orthogonally to ATP synthase oligomerization to modulate IMM morphology but is only required when PL saturation is increased.

### Modeling predicts compensatory roles for lipid and protein-encoded curvature in shaping cristae tubule formation

To understand the interaction between CL, which contributes to net membrane spontaneous curvature, and ATP synthase oligomers, whose induced curvature is localized specifically at cristae ridges, we employed a continuum modeling framework based on previous efforts to model membrane tubule formation (Mahapatra, 2022). We modeled the simplest CM structure — tubules — as axisymmetric tubes that bud from a flat membrane (Figure 5A). We added a pre-defined coat area of ATP synthase oligomers that contribute an anisotropic curvature (*D*_0_). An isotropic membrane curvature (*C*_0_) was then applied across the entire membrane to model the effects of asymmetrically localized CL. In simulations, we varied the magnitude of *D*_0_ and *C*_0_ based on data from simulations of curvature induced by ATP synthase dimers (Anselmi *et al*, 2018) and CL effects on mitochondrial membrane compositions estimated by simulations of outer and inner IMM leaflets (Figure EV3G). In both cases, the approximate ranges were set from 0 to 0.035 nm^-1^. The stiffness of the membrane was also varied across biologically reasonable values (10-20 k_B_T) and the membrane tension was set at 0.01 pN/nm (Hassinger *et al*, 2017). Finally, to incorporate the stresses due to the proposed roles of the MICOS complex and Mgm1p at the CJs, we used a localized collar force density of 8 pN/nm around the base of the tubule neck. Additional details of the governing equations and parameters of the model can be found in the Appendix.

**Figure 5:**
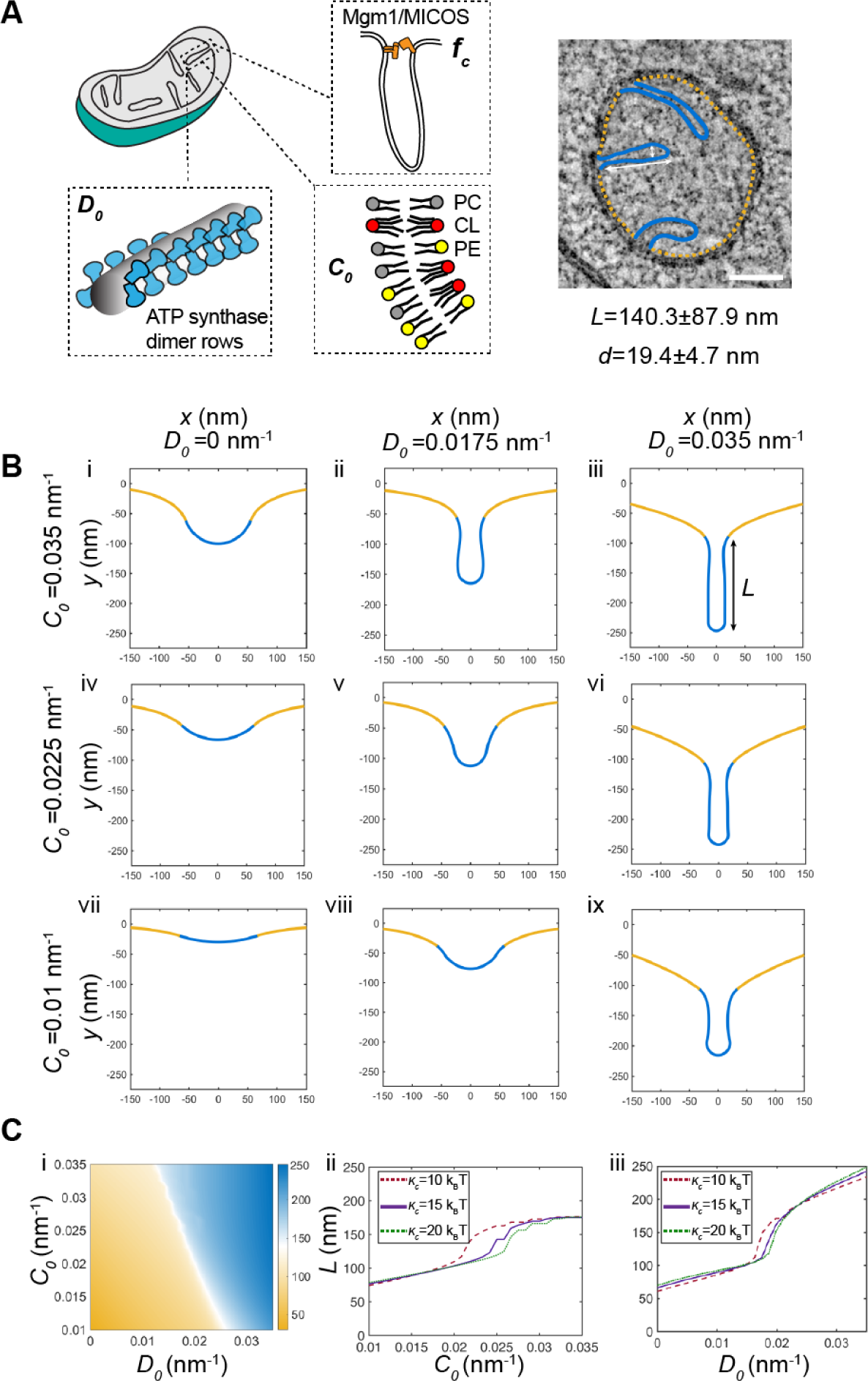
Continuum modeling of CM tubule formation reveals a snapthrough instability mediated by both lipid and ATP synthase generated curvatures. **(A)** Schematic depiction of yeast CM tubules containing a modeled neck force (*f*_c_) induced by Mgm1p and MICOS complexes, a deviatoric curvature imposed by ATP synthase at tubule ridges (*D*_0_), and a spontaneous curvature along the entire membrane imposed by asymmetric distribution of CL across the IMM (*C_0_*). (Right) TEM image showing typical yeast CM tubules in a single mitochondrion. Scale bars, 100 nm. The average tubule length (140.3±87.9nm) and diameter (19.4±4.7nm) of 15 cristae tubules were analyzed from tomograms of n=3 SFA2 mitochondria. **(B)** Changes in deviatory and spontaneous curvatures modulates tubule morphology. Panels i-ix show shapes of the membranes from simulations of the continuum model. For these simulations, the bending modulus of the membrane was maintained at 15 k_B_T and the tension was set at 0.01 pN/nm. The total membrane area was set to 5.65 x 10^5^ nm^2^ and the coated area was 1.413 x 10^4^ nm^2^. The values of *C_0_* and *D_0_* were varied as shown and *f*_c_ was set to 8 pN/nm at the base of the coat. **(C)** CM tubule formation shows a snap-through behavior. (i) Length of the tube as a function of *C_0_* and *D_0_*for the same values of bending modulus, tension, and areas as shown in (B). The white region shows the transition from the short to the long tube. The color bar shows the length of the tube in nm. Line graphs in (ii) show the length of the tube as a function of *C_0_* for *D_0_* fixed at 0.0175 nm^-1^ and line graphs in (iii) show the length of the tube as a function of *D_0_* for *C_0_* fixed at 0.0225 nm^-1^ for three different values of *к*. In both these graphs, the abrupt change in length is indicative of a snapthrough instability.

The model suggested that combination of *D*_0_ and *C*_0_ was sufficient to deform the membrane into different shapes reminiscent of flat membranes, buds that could be relevant for onion-like IMM formation, and CM-like tubules (Figure 5B). When tubules formed, their length (*L*) and diameter (*d*) were generally consistent with that of CMs in cells (Figure 5B). High values of *D*_0_ (Figure 5B, third column) promoted the formation of tubules for all values of *C*_0_ and bending moduli, and did so independently of the presence of the collar force (Appendix Figure S6). Low values of *D*_0_, mimicking a loss of ATP synthase dimers in *atp20*Δ or SFA3/4 cells, did not allow for tubule formation and instead led to flat or bud-like structures (Figure 5B, i-iii). The latter was observed even when *C*_0_, potentially encoded by CL, remained high. For intermediate values of *D*_0_, which could approximate the state in SFA2 cells that show partial loss of ATP synthase oligomerization, higher values of *C*_0_ were required to form tubules (Figure 5B, ii and v), while lower values resulted in shallow invaginations. This dependence on isotropic spontaneous curvature when anisotropic spontaneous curvature is partially lost could explain the interaction experimentally observed between CL and PL saturation. We note that similar calculations in the absence of the collar force (Appendix Figure S6), show bud-like and tubular structures with a wider neck, consistent with the model that local forces exerted by the MICOS and Mgm1p shape CJs.

Further investigation into the parameter space for cristae-like tubular structures revealed a transition from the short (low *L*) U-shaped invaginations to long (high *L*) tubules in an abrupt manner (Figure 5Ci). This transition was observed both as *D*_0_ (e.g. via ATP synthase oligomerization) increased for a fixed value of *C*_0_ (Figure 5Cii) or as *C*_0_ (e.g. via CL-encoded membrane spontaneous) was increased for a fixed value of *D*_0_ (Figure 5Cii) for the different values of bending moduli (see also Appendix Figure S6 for additional simulations). Such sudden changes to morphology, e.g. from a bud to a tube, in mechanical parameter space is termed a snapthrough instability or buckling event (Walani *et al*, 2015). In our model, the exact *C*_0_ and *D*_0_ values at which the snapthrough occurred was also modulated by the presence of a collar force at the tubule neck, suggesting that mechanics could underlie the functional interactions observed between Mic60p at the CJ and PL saturation (Figure EV2B, Appendix Figure S6).

### CL is an essential mitochondrial lipid in low oxygen environments that promote saturated lipidomes

The lack of phenotype for *crd1*Δ strains under standard laboratory growth conditions have long been perplexing, given the demonstrated importance of CL for mitochondrial function in other organisms (Paradies *et al*, 2014). Natural yeast environments, such as rotting fruits or fermentation tanks, are intrinsically microaerobic and inhibit Ole1p, which like other desaturases requires oxygen as a final electron acceptor. Oxygen binding to lipid desaturases is dependent on a low-affinity di-iron site (K_M_ ∼60 μM) and so desaturase activity is sensitive to environmental oxygenation (Kwast *et al*, 1999; Vasconcelles *et al*, 2001). Yeast grown in microaerobic fermenters show a lower level of di-unsaturated PLs than those grown under highly aerated conditions (low volume shake flasks), with a lipidome that was intermediate between that of aerated SFA2 and SFA3 cells (Figure EV5A, Figure EV5B). We hypothesized that in these less aerated environments, CL metabolism would have evolved to support an essential mechanical function.

To test the role of oxygenation on CL function, we grew *CRD1* and *crd1*Δ yeast strains in microaerobic chambers, comparing their lipidomes and mitochondrial morphologies to those grown in highly aerated shake flasks (Figure 6A). Under limited oxygenation, *CRD1* cells increased the abundance of CL twofold compared to highly aerated conditions (Figure 6B) and showed increased staining by the CL-binding dye nonyl acridine orange (NAO) (EV5C). This increase did not accompany any general changes to mitochondrial volume or length in aerobic vs. microaerobic cells (Figure EV5D). *CRD1* and *crd1*Δ cells showed identical increases in the saturation of PLs under microaerobic conditions (Figure 6C, Figure EV5B) and both retained ATP synthase dimers (Figure EV5E). After microaerobic growth, yeast containing CL still presented tubular mitochondrial morphologies, but *crd1*Δ cells predominantly displayed aberrant mitochondrial morphologies (Figure 6D-E). Ultrastructural analysis of IMM structure revealed that *CRD1* cells grown under microaerobic conditions displayed tubular cristae morphologies while *crd1*Δ cells displayed a mixture of flat and onion-like IMM structures (Figure 6F). CL can thus be required for CMs in yeast, but not in highly oxygenated laboratory conditions that suppress PL saturation.

**Figure 6:**
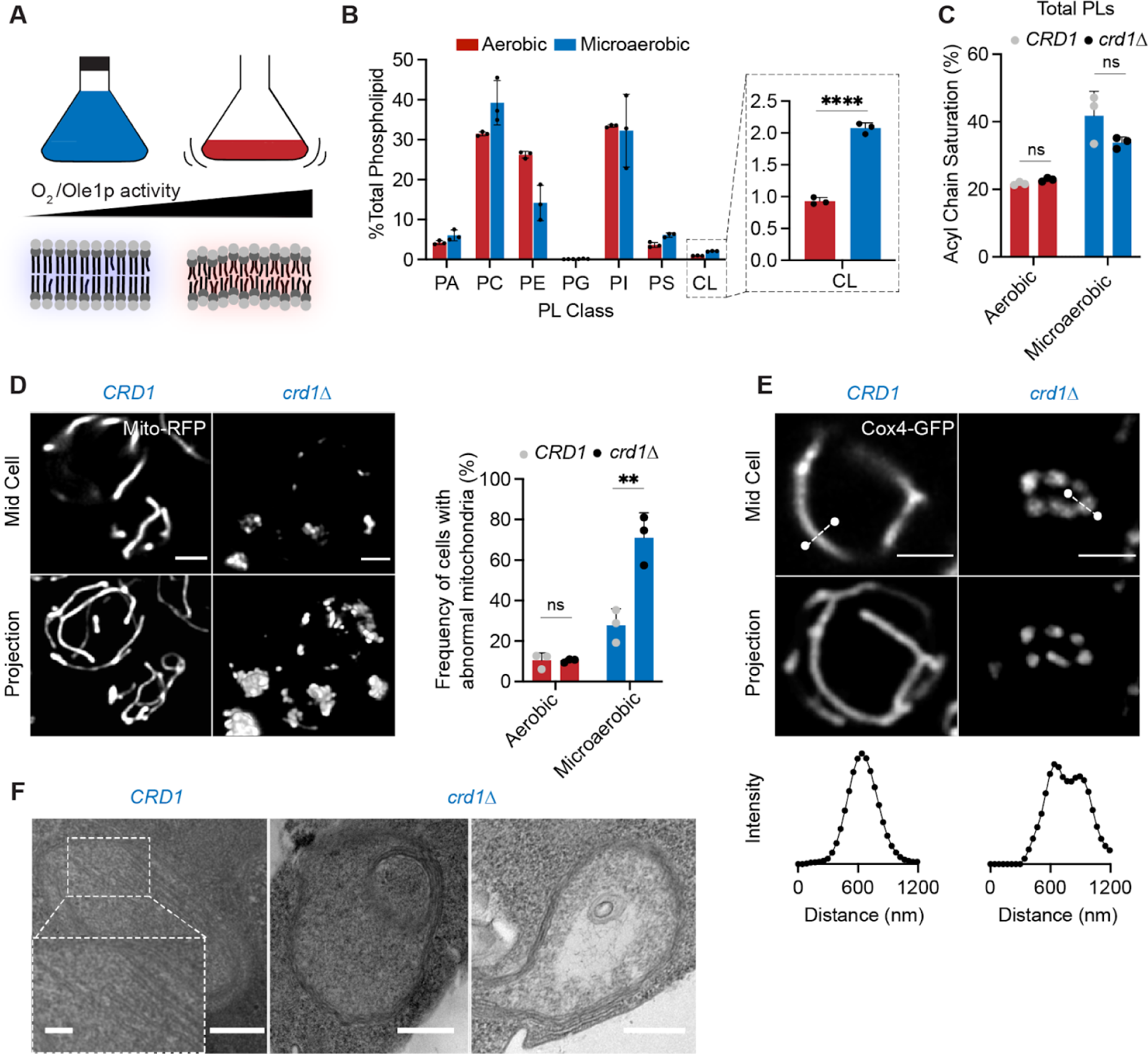
Cardiolipin synthesis is essential for mitochondria in yeast growth conditions characterized by low oxygenation. **(A)** Schematic representation of different oxygen concentrations in different yeast growth environments. Microaerobic conditions cause increased saturation of membranes due to lower desaturate activity of Ole1p, an oxygen dependent enzyme. **(B)** WT cells increase the abundance of CL under microaerobic conditions. Shown are abundances of each PL class, n=3 biological replicates. Error bars indicate SD. ****p < 0.0001, unpaired two-tailed t-test compared against wild-type. **(C)** Microaerobic growth conditions cause an increase in acyl chain saturation in the total PL pool, which is not affected by loss of CL in *crd1*Δ strains. Error bars indicate SD from n=3 biological replicates. **(D)** Under microaerobic conditions, loss of CL (*crd1*Δ) causes loss of tubular mitochondrial structure. WT (*CRD1*) and *crd1*Δ cells were grown in microaerobic chambers for 48 hours prior to imaging, n=3 biological replicates. Scale bars, 2 μm. **p < 0.005, unpaired two-tailed t-test compared against WT. **(E)** Microaerobic *crd1*Δ show hollow mitochondria, imaged with Cox4-GFP. Scale bars, 2 μm. Line profile analysis (below) depicts fluorescent intensity across the indicated mitochondria. **(F)** Under microaerobic conditions, *CRD1* cells contain long, sheet-like cristae structures while *crd1*Δ cells lack cristae, and display both onion-like and flat abnormal IMM structures as visualized by thin-section TEM. Scale bars, 250 nm. Inset showing abundant cristae sheets in *CRD1* cells, scale bar, 125 nm.

## Discussion

In this study, we explored an initial observation that small changes to the composition of yeast PLs cause the loss of CMs in mitochondria. We found that saturated PLs inhibit ATP synthase oligomerization, which is progressively lost when either the expression or activity of the lipid desaturase Ole1p decreases. Based on molecular modeling of IMM lipidomes, we hypothesized that conical lipids buffer against loss of ATP synthase as a cristae-shaping protein. We explored the interaction between the curvature at cristae ridges induced by ATP synthase oligomerization and modulated by PL saturation, and lipid-encoded curvature across the IMM, which is promoted by conical lipids like CL. Both factors contribute to whether the IMM exists in a high or low curvature state, which in turns controls ATP generation and mitochondrial fitness. The sharp transition we initially observed between these states can be understood through a continuum model that highlights a snapthrough instability, a phenomenon in which a system can shift between two morphological states via modest perturbations in its material properties or forces applied. Also termed mechanical buckling, such events are intrinsic to membrane bending mechanics (Ou-Yang *et al*, 1999; Vasan *et al*, 2020) and have been proposed to act in other curved cellular structures, such as endocytic buds (Walani *et al*, 2015; Hassinger *et al*, 2017).

Based on our observations, we propose that the intrinsic membrane curvature needed to form CMs can be generated both by cristae-shaping proteins and by asymmetric distributions of lipids across the bilayer, which contributes net spontaneous curvature between the two leaflets. CL is a negatively curved PL in the IMM for which there are multiple potential sources of asymmetry in the IMM. Staining with NAO suggests that the yeast IMM features up to 2-fold more CL in the outer leaflet (Petit *et al*, 1994; Gallet *et al*, 1997), which would optimize its negative curvature at cristae ridges. The differences in local pH between the matrix and IMS could also favor the monoanion CL species in the IMS-facing leaflet, which would also promote negative curvature at cristae ridges. Finally, CL and other conical lipids segregate into deformed membrane regions, further promoting curvature. Such a phenomenon has been observed experimentally when CL localized to thin, high curvature tubules that have been pulled from large, low curvature vesicles (Beltrán-Heredia *et al*, 2019). In simulations, this effect is apparent in localized CL concentrations in thermal fluctuations (Figure 3G) and those induced by compression (Dahlberg & Maliniak, 2010; Boyd *et al*, 2017). In the IMM, it is likely that local deformations induced by ATP synthase (Blum *et al*, 2019) cause the local concentration of conical lipids (e.g. CL and di-unsaturated PE), which further act to deform the IMM. The action of membrane shaping lipids and proteins are therefore likely to be intrinsically linked.

The interplay of CL with PL saturation provides a biophysical rationale for long-standing questions regarding its role in the IMM. The consequences of CL loss, widely explored to understand the pathophysiology of Barth syndrome, differ strikingly across experimental systems. CL is essential in both mice (Kasahara *et al*, 2020; Xu *et al*, 2021) and mammalian cell lines (Choi *et al*, 2007), but not in yeast or *Drosophila*, where its loss has more subtle effects on respiration (Xu *et al*, 2021). The mechanical functions of CL function could be dependent on the surrounding lipid environment in the IMM, which can change depending on growth conditions. Oxygenation is one natural modulator of PL saturation due to the intrinsic enzymology of lipid desaturases. In yeast, growth under microaerobic conditions leads to lipidomes with predominantly monounsaturated PLs that require CL to generate CMs in the IMM. In contrast, growth under highly aerated laboratory conditions show an unusually high level of di-unsaturated PLs (Appendix Figure S1G) and does not necessitate CL.

The role of oxygenation in controlling PL saturation is not exclusive to yeast. In cancer cells, hypoxia reduces activity of SCD1 (stearoyl CoA desaturase 1), which also drives lipidome remodeling (Kamphorst *et al*, 2013; Ackerman *et al*, 2018). Specific tissue environments, such as in the gut, are also microaerobic (Zheng *et al*, 2015), and a recent genome-wide screen implicated CL metabolism in low oxygen fitness of intestinal T-cells (Reina-Campos *et al*, 2023). In human embryonic kidney (HEK) 293 cells, the morphological and respiratory effects of knocking down cardiolipin synthase (*CRLS1*) are potentiated by increased saturation caused by either mild PA treatment or low oxygen growth (Figure EV6). In PA treated *CRLS1* knockdown cells, the density and length of cristae sheets are strongly reduced (Figure EV6D-6E). Thus, the interaction between CL and PL saturation is likely to extend beyond yeast mitochondria.

An outstanding question of this work is how specific lipid perturbations modulate the dimerization and higher order organization of ATP synthases. PL saturation, which influences both membrane stiffness and curvature, is a unique regulator of ATP synthase organization in cells: yeast strains that have drastically altered mitochondrial PC, PE and CL levels all retain dimers (Claypool *et al*, 2008; Baile *et al*, 2014; Baker *et al*, 2016). Among membrane protein dimers, the yeast ER sensors Mga2p/Spt23p show a similar dimer to monomer transition when their host membrane shifts di-unsaturated to mono-unsaturated PLs (Covino *et al*, 2016; Ballweg *et al*, 2020). While Mga2p/Spt23p are small, single pass transmembrane proteins, the CLC Cl^-^/H^+^ antiporter dimer has a similar buried surface area as ATP synthase (∼2400 Å^2^) and a modest dimerization free energy (19 k_B_T) (Chadda *et al*, 2016) that is also highly sensitive to its lipidic environment (Chadda *et al*, 2021). In CLC, short chain PLs have been shown to stabilize the monomer conformation and through this mechanism weaken its dimerization. Unlike CLC, dimerization of ATP synthase is associated with an extreme local membrane deformation – a ∼90° bend in *S. cerevisiae* (Guo et al, 2017). One hypothesis is that the changes in mechanical properties encoded by PL saturation, especially stiffness, mediate local membrane deformation, reducing the dimer:monomer equilibrium by increasing its elastic cost. Distinguishing between these models will require future experiments and modeling, which would be aided by structures of monomeric ATP synthases with resolved dimerization subunits.

During the evolution of eukaryotic cells, the machinery for ATP generation in the ETC adopted a secondary function in the shaping of the IMM. The emergence of ATP synthase dimers and other cristae-shaping protein complexes occurred alongside a specialization in the inner membrane lipidome, including the proliferation of CL from the proteobacterial inner membrane (Sohlenkamp & Geiger, 2016; Rowlett *et al*, 2017). Among extant eukaryotes, ATP synthase organization varies widely; for example, in mammalian mitochondria ATP synthases range from primarily monomeric (Galber *et al*, 2019) to mixtures of monomers and dimers (Bisetto *et al*, 2007). It is notable that CL is essential for proper IMM structure in these systems, unlike in aerobic yeast in which dimers are predominant. The composition of other cristae-shaping proteins, such as the MICOS complex (Huynen *et al*, 2016), also differ across eukaryotes, despite the ubiquity of CMs in mitochondria. Such variability in IMM shaping proteins could have necessitated the utilization of alternative forms of curvature generation encoded by the mitochondrial lipidome.

## Materials and Methods

### Strains and growth media

The yeast strains used in this study can be found in Appendix Table S3. Yeast cells were grown in YPEG (1% Bacto yeast extract, 2% Bacto peptone, 2% Ethanol, 2.5% Glycerol), YPD medium (1% Bacto yeast extract, 2% Bacto peptone, 2% glucose) or complete supplement mixture (CSM, 0.5% Ammonium Sulfate, 0.17% yeast nitrogen base without amino acids and 2% glucose) lacking appropriate amino acids for selection. Yeast mutants were generated by PCR-based homologous recombination, ORFs of the gene of interest were replaced by either *KanMX* or *His3MX6* cassettes. For *OLE1* promoter substitution, a set of previously generated mutant *TEF1* promoters were utilized (Alper *et al*, 2005). An additional, weaker promoter (*Pm1)* was generated by error prone PCR and added to this set. Promoter substitution was completed on the haploid base strain W303a as previously described (Degreif *et al*, 2017).

### Yeast physiology

Growth on non-fermentable carbon sources were assayed using growth curves on 24-well plates (Avantor) sealed with a gas permeable film (Diversified Biotech). Cells were first grown in biological triplicate overnight in complete synthetic medium (CSM) containing 2% glucose. Cells were back-diluted 1:100 in fresh CSM containing 2% glucose or 3% glycerol and shaken in a plate reader (Tecan) for 48 hours. Specific growth rates were extracted from the exponential phase of growth. For viability assays, cells were first grown in the same fashion as for yeast growth curves but were serially diluted onto CSM plates (1:5 successive dilutions) containing either 2% glucose or 3% glycerol and 2% ethanol. Plates were grown for 3 days on glucose plates, and for 4 days for ethanol/glycerol plates.

For microaerobic growth, cells were pre-incubated overnight in CSM without uracil (2% glucose) in a controlled temperature shaker at 30°C. Cells were then back diluted into fresh synthetic medium and grown to stationary phase in home-built microaerobic chambers with limited oxygen supply. Chambers consisted of glass culture tubes with tight fitting rubber caps; tubing allowed for gas efflux into an attached bubbler. Live cell imaging was conducted on aliquots, and cell pellets were resuspended in sterile water, lysed and flash frozen for lipidomics analysis.

### Lipidomics

Lipid compositions of whole cells and isolated mitochondria from yeast strains were conducted at Lipotype GmbH (Dresden, Germany). Mass spectrometry-based lipid analysis was performed as previously described (Ejsing *et al*, 2009; Klose *et al*, 2012). Lipids were extracted using a two-step chloroform/methanol procedure (Ejsing *et al*, 2009). Samples were spiked with an internal lipid standard mixture containing: CL 14:0/14:0/14:0/14:0, ceramide 18:1;2/17:0 (Cer), diacylglycerol 17:0/17:0 (DAG), lyso-phosphatidate 17:0 (LPA), lyso-phosphatidyl-choline 12:0 (LPC), lysophosphatidylethanolamine 17:1 (LPE), lyso-phosphatidylinositol 17:1 (LPI), lysophosphatidylserine 17:1 (LPS), phosphatidate 17:0/14:1 (PA), phosphatidylcholine 17:0/14:1 (PC), PE 17:0/14:1, PG 17:0/14:1, PI 17:0/14:1, phosphatidylserine 17:0/14:1 (PS), ergosterol ester 13:0 (EE) and triacylglycerol 17:0/17:0/17:0 (TAG). After extraction, the organic phase was transferred to an infusion plate and dried in a speed vacuum concentrator. 1st step dry extract was re-suspended in 7.5 mM ammonium acetate in chloroform/methanol/propanol (1:2:4, v:v:v) and 2nd step dry extract in 33% ethanol solution of methylamine in chloroform/methanol (0.003:5:1; v:v:v). All liquid handling steps were performed using Hamilton Robotics STARlet robotic platform with the Anti Droplet Control feature for organic solvents pipetting. Samples were analyzed by direct infusion on a QExactive mass spectrometer (Thermo Scientific) equipped with a TriVersa NanoMate ion source (Advion Biosciences). Samples were analyzed in both positive and negative ion modes with a resolution of R_m/z=200_=280000 for MS and R_m/z=200_=17500 for MSMS experiments, in a single acquisition. MSMS was triggered by an inclusion list encompassing corresponding MS mass ranges scanned in 1 Da increments (Surma *et al*, 2015). Both MS and MSMS data were combined to monitor EE, DAG and TAG ions as ammonium adducts; PC as an acetate adduct; and CL, PA, PE, PG, PI and PS as deprotonated anions. MS only was used to monitor LPA, LPE, LPI and LPS as deprotonated anions; Cer and LPC as acetate adducts. Data were analyzed with in-house developed lipid identification software based on LipidXplorer (Herzog *et al*, 2011, 2012). Data post-processing and normalization were performed using an in-house developed data management system. Only lipid identifications with a signal-to-noise ratio >5, and a signal intensity 5-fold higher than in corresponding blank samples were considered for further data analysis.

The acyl chain composition of HEK293 cells was analyzed by Bligh-Dyer extraction followed by Gas Chromatography coupled to Mass Spectrometry (GC-MS) of transesterified fatty acid methyl esters (FAME) on an Agilent 8890-5977B GC-MS, as previously described (Winnikoff *et al*, 2021). Quantification was performed in Agilent MassHunter using external FAME standards (Cayman Chemical #20503).

### Mitochondria purification

Yeast mitochondria were isolated from 1L of yeast cells grown in YPEG, (1% Bacto yeast extract, 2% Bacto peptone, 2% Ethanol, 3% Glycerol) YPD medium (1% Bacto yeast extract, 2% Bacto peptone, 2% glucose) or CSM with 2% glucose (microaerobic conditions) at 30°C as previously described (Gregg *et al*, 2009; Meisinger *et al*). Cells were grown to stationary phase and harvested in a buffer consisting of 100mM Tris/H_2_SO_4_ (pH 9.4) and 10mM dithiothreitol. Spheroplasts were formed from digestion of the cell wall using Zymolyase 20-T (MP Biomedicals) in a buffer containing 20mM Potassium Phosphate (pH 7.4) and 1.2M Sorbitol. Spheroplasts were lysed by homogenization using a glass homogenizer and subsequently centrifuged to remove unbroken cells, large debris, and nuclei. Enriched mitochondria were pelleted and resuspended in SEM buffer (10mM MOPS/KOH (pH 7.2), 250mM Sucrose and 1mM EDTA), snap frozen and stored for up to 1 month at −80 °C. To obtain purified mitochondria, bereft of contamination from other organelles, such as microsomes and vacuoles, the crude mitochondrial fraction was subjected to sucrose density cushion ultracentrifugation. Cushions are poured containing 60% and 32% (w/v) sucrose concentrations in EM buffer (10mM MOPS/KOH (pH 7.2), 1mM EDTA). Density cushions containing crude mitochondrial samples in SEM buffer are subjected to ultracentrifugation in a SW32 Ti swinging-bucket rotor for 1 hour at 100,000 x g at 4 °C. The yellow/brown band at the 60/32% (w/v) sucrose interface is removed and centrifuged to pellet purified mitochondria for subsequent analysis. Mitochondrial protein quantity was determined via BCA assay.

Mitoplasts containing an intact inner mitochondrial membrane stripped of the outer membrane were isolated as previously described (Zhang *et al*, 2005). After initial incubation of isolated mitochondria in a hypotonic buffer followed by centrifugation for 10 minutes at 14,000 x g at 4 °C. Mitoplasts were resuspended in SEM buffer prior to lipidomic analysis.

### Respirometry

Oxygen consumption rates (OCR) of whole cell yeast strains were measured with a Clark electrode (YSI 5300A Biological Oxygen Monitor System). Cells were grown in biological replicates overnight in CSM containing 0.4% glucose (starvation conditions). Cells were then back-diluted into fresh CSM and grown to OD 0.4-0.6. OCRs were quantified after initial stirring of culture for 3 minutes in a thermostatically controlled chamber at 30 °C. Respiration rates were determined after normalization to OD of the sample. Respirometry on HEK293 cells were performed on a SeaHorse XF Pro (Agilent) using the real-time ATP rate assay kit (103591-100) on 10,000 cells per replicate per condition.

### Live cell microscopy

All live cell microscopy was conducted using Plan-Apochromat 63x/1.4 Oil DIC M27 on the Zeiss LSM 880 with Airyscan (default processing settings); image acquisition and processing were performed using ZEN software. In yeast experiments, cells were plated on 8 well chambered coverglass (Nunc Lab-Tek) pre-incubated with concanavalin A (MP Biomedicals). To assess mitochondrial morphology in yeast, cells were initially grown in biological replicates overnight in CSM containing 0.4% glucose under selection conditions. Cells were then back diluted and imaged in an exponential phase. For analysis, yeast cells were split into 3 groups based on mitochondrial morphology: ‘aberrant’ groups display a punctate cluster at the center of the cell, whilst ‘fragmented’ groups display a lack of tubular morphology and consist of individual mitochondrial puncta separated within the cell (Appendix Figure S2). Normal mitochondrial morphology is associated with tubes spanning the length of the cell that display regular lateral motion. Percentage ‘mitochondrial abnormality’ is derived from combining the amount of cells with an aberrant or fragmented morphology divided by the total number of cells. At least 50 cells were quantified in each sample and replicates were of individually grown cultures. To visualize the yeast cell wall, cells were stained with calcofluor white (Sigma-Aldrich), which binds chitin within the cell wall. Mitochondrial volume and length was quantified using Mitograph software as previously described (Viana *et al*, 2015). Analysis of HEK293 mitochondria was done by staining cells grown on glass bottom dishes (MatTek) with MitoTracker Deep Red (ThermoFisher Scientific M22426) for 30 minutes prior to imaging.

Membrane potential was assayed by growing yeast to exponential phase in CSM containing 0.4% glucose followed by staining with 200 nM TMRE (Thermo Fisher Scientific T669) for 20 minutes. Cells were then washed twice with water prior to imaging. For uncoupled conditions, cells were incubated with 20 μM carbonyl cyanide m-chlorophenyl hydrazone (CCCP, Sigma-Aldrich) for 10 minutes prior to incubation with TMRE. For analysis of membrane potential in ATP synthase inhibited conditions, cells were incubated with 5 μM oligomycin for 45 minutes prior to incubation with TMRE. To analyze yeast mtDNA nucleoids, cells were grown to exponential phase in CSM containing 0.4% glucose and washed once with PBS prior to staining with SYBR Green I (SGI, Thermo Fisher Scientific S7563) for 10 minutes. Cells were then washed three times with PBS prior to imaging. Relative yeast CL content in aerobic and microaerobic growth conditions was determined by staining cells with NAO (Thermo Fisher Scientific A1372). Cells were stained with 100nM NAO for 20 minutes and then washed three times with water prior to imaging. The maximum intensity per cell was determined using profile analysis in ImageJ.

### Blue native-PAGE analysis

Isolated mitochondria solubilized in digitonin (0.5:1 g/g protein) were assayed by BN-PAGE as previously described (Timón-Gómez *et al*, 2020), with minor modifications. 200-400 μg of mitochondria were incubated with digitonin for 10 minutes prior to centrifugation at 20,000 x g for 30 mins at 4 °C. The subsequent supernatant was mixed with native PAGE buffer and glycerol and loaded onto precast native PAGE gels (Invitrogen). ATP synthase dimerization state was probed using an anti-ATP synthase primary antibody (Rak & Tzagoloff, 2009) (1:1000) and anti-rabbit IgG secondary antibody (Thermo Fisher Scientific). Supercomplex formation was assessed using an anti-Cox1p (CIV) antibody and anti-mouse secondary antibody (Thermo Fisher Scientific).

### Immunoblot analysis

Whole cell yeast lysates were grown in YPEG and 2.5 OD units were subjected to protein extraction and SDS-PAGE as previously described (Kushnirov, 2000). After transfer to PVDF membranes, western blot analysis was completed using the following primary antibodies at stipulated dilutions in blocking buffer (5% BSA in TBST): 1:1000 for Cox4p and Dpm1p and 1:250 for Pho8p. For analysis of isolated mitochondria, 10μg of total protein was loaded on SDS-PAGE gels and transferred to PVDF membranes prior to western blotting with the aforementioned antibodies as well as with the Mgm1p and Mic60p antibodies (Rabl *et al*, 2009). In HEK293 cells, siCRLS1 knockdowns were verified by immunoblotting with anti-CRLS1 polyclonal antibody and a polyclonal Actin loading control. All antibodies are listed in Appendix Table S5.

### Electron microscopy

Blocks of late exponential phase yeast cells were prepared either by high-pressure freezing/freeze substitution (HPF-FS) (McDonald & Müller-Reichert, 2002) (SFA4, SFA2*crd1*Δ, *atp20*Δ) or chemical fixation followed by partial cell wall digestion (Bauer *et al*, 2001) (SFA2, SFA4, *crd1*Δ and microaerobic cells) as previously described. Cell wall digestion was required for high contrast staining of WT-like tubular CM for 3D segmentation. For microaerobic cells, chemical digestion was performed with 0.25 mg/mL zymolyase-20T for one hour at room temperature. Aerobically-grown mutant cells lacking CMs showed sufficient membrane contrast in HP-FS samples. Thin sections about 60 nm thick were cut from the blocks of yeast with a Leica ultramicrotome and placed on 200-mesh uncoated thin-bar copper grids. A Tecnai Spirit (FEI, Hillsboro, Oregon) electron microscope operated at 120 kV was used to record images with a Gatan Ultrascan 4K x 4K CCD camera at 6.0, 2.9, and 1.9 nm/pixel. For TEM on HEK 293 mitochondria, cells were grown to confluency in MatTek dishes coated with fibronectin, prepared and recorded as previously described (Darshi *et al*, 2011).

### Tomography

Semi-thick sections of thickness about 300 nm were cut from the blocks of yeast with a Leica ultramicrotome and placed on 200-mesh uncoated thin-bar copper grids. 20-nm colloidal gold particles were deposited on each side of the grid to serve as fiducial cues. The specimens were irradiated for about 20 min to limit anisotropic specimen thinning during image collection at the magnification used to collect the tilt series before initiating a dual-axis tilt series. During data collection, the illumination was held to near parallel beam conditions and the beam intensity was kept constant. Tilt series were captured using SerialEM (University of Colorado, Boulder) software on a Tecnai HiBase Titan (FEI) electron microscope operated at 300 kV and 0.81 nm/pixel. Images were recorded with a Gatan 4K x 4K CCD camera. Each dual-axis tilt series consisted of first collecting 121 images taken at 1 degree increment over a range of −60 to +60 degrees followed by rotating the grid 90 degrees and collecting another 121 images with the same tilt increment. To improve the signal-to-noise ratio, 2x binning was performed on each image by averaging a 2×2 x-y pixel box into 1 pixel using the newstack command in IMOD (University of Colorado, Boulder). The IMOD package was used for tilt-series alignment, reconstruction, and volume segmentation. R-weighted back projection was used to generate the reconstructions.

### Mesh generation and analysis

3D *in silico* reconstructions of mitochondria were generated from electron-tomographic images. The software IMOD was used to trace the mitochondrial membranes in 2D: the outer leaflet of the OM, and the inner leaflet of the IBM and CM were manually traced as separate objects, following procedures previously described (Mendelsohn *et al*, 2022). Subsequently, 2D traces were imported into Blender using the NeuropilTools module in CellBlender. The program Contour Tiler (Edwards *et al*, 2014) — integrated with NeuropilTools — was used to generate 3D triangulated meshes in Blender. The triangulation was performed individually for each membrane object in each mitochondrion. Afterwards, the Boolean Difference Modifier was used to subtract the CM object from the IBM object, generating in this manner the CJs in the IBM. The meshes were refined with the Smooth and Normal Smooth improvement tools from GAMer2 (Lee *et al*, 2020). Curvature calculations were carried out with GAMer2, using the MDSB algorithm. For all curvature analysis, the smooth curvature after one iteration was considered. This smoothing represents the average curvature of a vertex and its neighbors. Surface areas and volumes were calculated using the CellBlender add-on in Blender.

### Mammalian cell culture

HEK293 cells (Sigma Aldrich) and were cultured in DMEM (Gibco) supplemented with 10% FBS (Gibco) at 37 °C in humidified air containing 5% CO_2_. For silencing experiments, Lipofectamine RNAiMAX was used per manufactures’ instructions for an siRNA final concentration of 10 nM, and cells were imaged or treated 24 hours after transfection. The siRNA constructs (ThermoFisher Scientific) included a non-targeting control (Catalog #4390843) and previously validated *CRLS1*-targeting sequence (siRNA ID: s29306) (Ohlig *et al*, 2018; Yang *et al*, 2023). For PA treatment, cells were transfected into complete DMEM containing specified 50 μM PA complexed to BSA and analyzed 24 hours after transfection. For aerobic/microaerobic incubations, media was replaced with DMEM containing delipidated FBS before incubation in either normoxic or microaerobic conditions 24 hours after transfection; the latter was maintained by continual flushing with nitrogen as previously described (Doedens *et al*, 2013). Microaerobic grown cells (1% oxygen) were incubated for 72 hours and aerobic grown cells (21% oxygen) were incubated for 48 hours before analysis to allow for two doublings each.

## Data Availability

All strains and plasmids are available upon request to the corresponding author. This study includes no data deposited in external repositories.

## Acknowledgments

José Faraldo-Gómez, Edward Lyman, Nicolas-Frédéric Lipp and Yi-Ting Tsai provided helpful discussions. The Herzik lab provided biochemistry assistance. Daniel Degreif and Sterling Ramsey assisted in strain development. Miguel Reina-Campos assisted with microaerobic cell culture experiments. Alexander Tzagoloff, Leticia Franco, Mário Barros and Andreas Reichert supplied antibodies. The University of California, San Diego - Cellular and Molecular Medicine Electron Microscopy Core (UCSD-CMM-EM Core, RRID: SCR_022039) provided equipment access and technical assistance. The UCSD-CMM-EM Core is partly supported by the National Institutes of Health Award number S10OD023527. The National Institutes of Health (R35-GM142960 to I.B, R01AG065549, R24GM137200 and U24NS120055 to M.H.E.), the Office of Naval Research (ONR N00014-20-1-2469 to P.R.), the National Science Foundation (DBI-2014862 to M.H.E), the Department of Energy (DE-SC0022954 to I.B.) and the Moore–Simons Project on the Origin of the Eukaryotic Cell (GBMF-9734 to I.B., P.R., and M.H.E.) provided financial support. K.V. was supported by the NIH Molecular Biophysics Training Grant (T32-GM008326C). C.T.L. was supported by a Kavli Institute for Brain and Mind Postdoctoral Fellowship. Howard Hughes Medical Institute supported initial project conception through the Janelia Visiting Scientist Program. Molecular dynamics simulations were run on hardware hosted by the Triton Shared Computing Cluster.

## Author Contributions

K.V., D.M., and I.B. designed and carried out the experiments. C.T.L., G.C.G., A.M., and P.R. designed and carried out the simulations. G.P., K-Y. K., H.A.P., S.P., and M.H.E. carried out electron microscopy. I.B and P.R. supervised the work. All authors contributed to data analysis and writing of the manuscript.

## Expanded View & Appendix

**Figure EV1:**
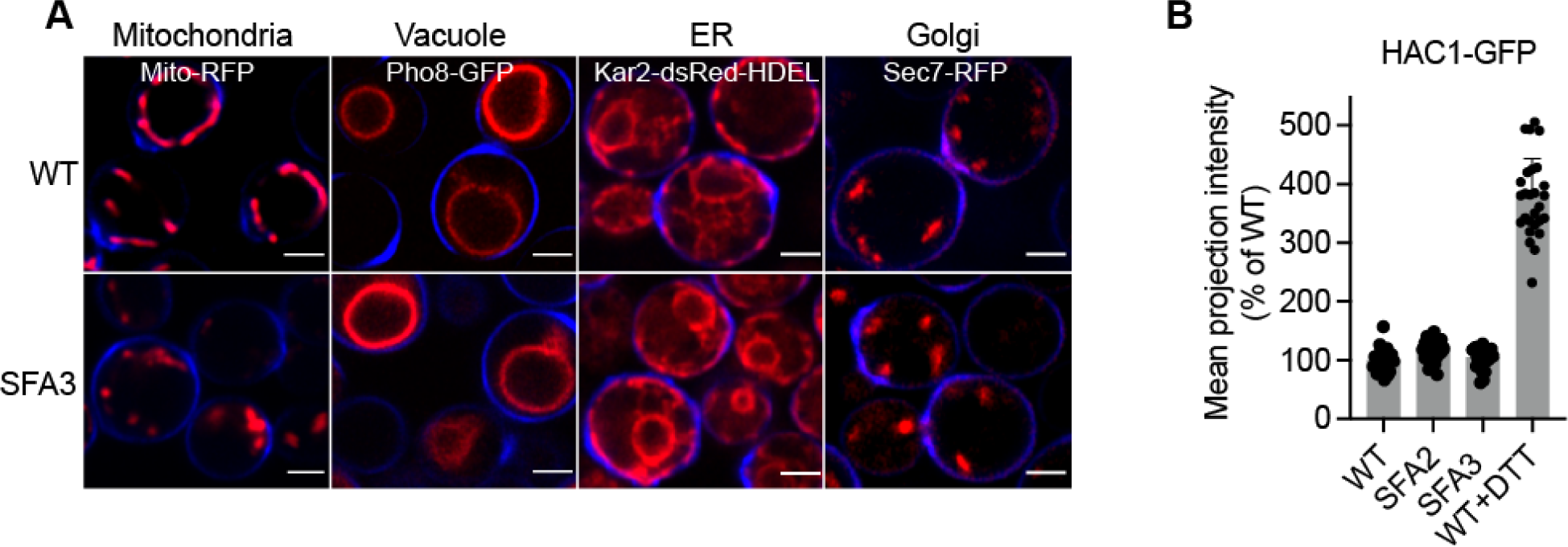
PL saturation causes defects to mitochondrial morphology while other organelles remain intact. (A) Decreasing expression of Ole1p expression results in abnormal mitochondria in SFA3 while other organelles remain intact. Organelles were imaged in cells that were transformed with plasmids expressing the following fusion proteins: mts-RFP (Mitochondria), Pho8-GFP (Vacuole), Kar2-dsRed-HDEL (ER), Sec7-RFP (Golgi). Cells were grown to exponential phase and were stained with cell wall-binding calcofluor white (blue). Scale bars, 2 μm. (B) ER stress was measured through induction of the unfolded protein response (UPR) using a HAC1-GFP splice reporter as previously described. GFP intensity was quantified in 3D projections from N=20 cells; the transition between SFA2/3 did not show any increase in UPR activation. WT cells treated with 2 mM dithiothreitol (DTT) were used as a positive control.

**Figure EV2:**
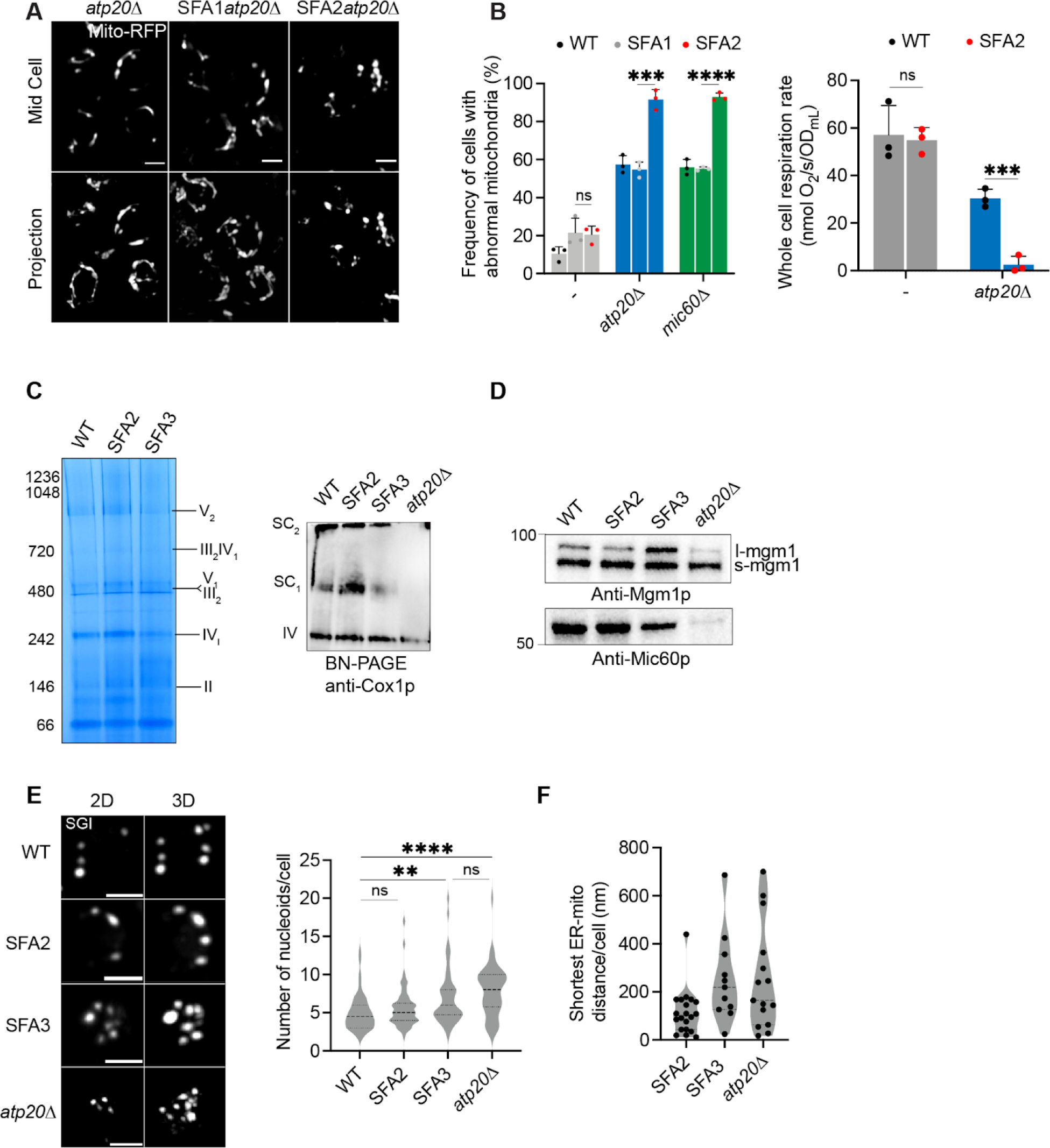
Interactions between cristae-shaping proteins, lipid saturation, and mitochondrial phenotypes. **(A)** Representative Airyscan confocal micrographs of yeast expressing matrix-localized RFP (mts-RFP). Scale bars, 2 μm. **(B)** (Left) Frequency of mitochondrial abnormalities as assayed by analysis of mts-RFP. N>50 cells were counted in biological triplicate (n=3) in each condition. Error bars indicate SD. ***p=0.0006, ****p <0.0001 unpaired two-tailed t-test compared SFA2*atp20*Δ and SFA2*mic60*Δ against SFA1*atp20*Δ and SFA1*mic60*Δ respectively. (Right) Respiration rates in *atp20*Δ and SFA2*atp20*Δ cells measured in biological triplicate (n=3) using a Clark electrode. Error bars represent SD. ***p < 0.0005, unpaired two-tailed t-test compared against *atp20*Δ. **(C)** SFA2/3 mitochondria do not show defects in ETC complexes or SC formation. BN-PAGE on digitonin-solubilized isolated mitochondria from SFA strains and *atp20*Δ revealed no changes in ETC complex levels (left) or supercomplexes (top right). Mitochondria from *atp20*Δ cells show defects in formation of Complex IV-containing supercomplexes. **(D)** Increasing lipid saturation does not change the status of non-ATP synthase cristae-shaping proteins. To determine the presence of Mgm1p and Mic60p in SFA strains, 10μg of isolated mitochondria were loaded and subjected to immunoblotting against anti-Mgm1p (C-term) and anti-Mic60p antibodies, respectively. **(E)** Both SFA3 and *atp20*Δ cells showed increased abundance of mtDNA nucleoids compared to WT. Nucleoids were visualized after staining with SYBR Green I (SGI) and the number of nucleoids per cell (N=50 cells) were quantified. Error bars indicate SD. Scale bars, 1 μm. ****p<0.0001, p=0.0020, unpaired two-tailed t-test against WT for *atp20*Δ and SFA3 respectively. **(F)** SFA3 and *atp20*Δ cells showed increased ER-mitochondria contact distances as analyzed by thin-section TEM. In each condition, the ER-mitochondria distances were measured using IMOD 3D software. Error bars represent SD. N=15 micrographs were quantified for each condition.

**Figure EV3:**
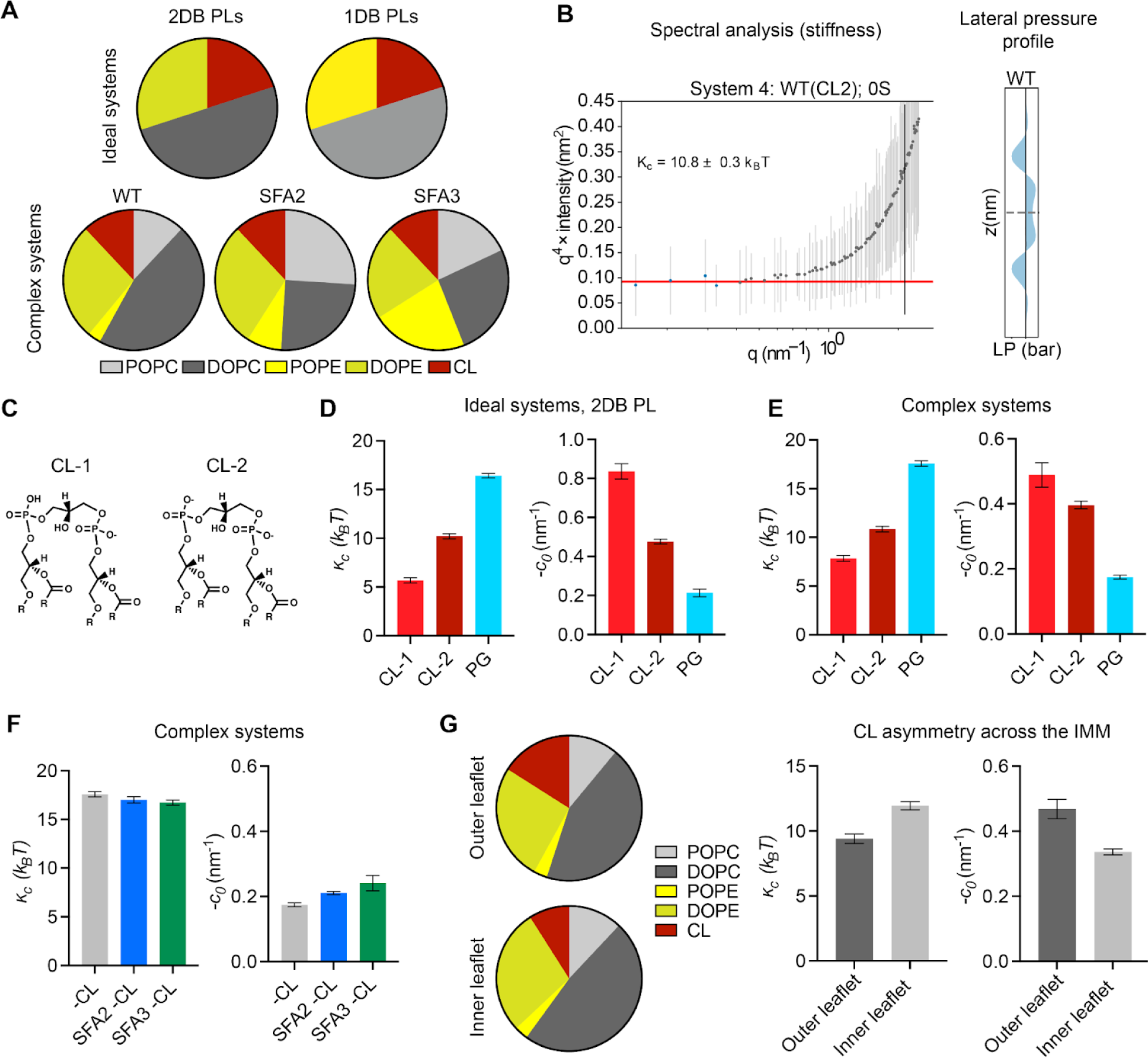
Coarse-grained molecular dynamics predicts an important role for CL in shaping membrane mechanical properties. **(A)** Ideal systems contained lipid compositions of fixed abundance, while changing the unsaturation of the acyl chains from di-unsaturated to monounsaturated. In contrast, the ‘complex’ systems mimicking the mitochondrial lipidomes of SFA strains accounted for headgroup adaptations to increasing saturation, such as increasing PE. Full compositions are listed in Appendix Table S2. **(B)** Example spectral analysis of thermal undulations, used to calculate bending moduli, and lateral pressure profiles, used to calculate spontaneous curvature. **(C)** Chemical structures of CL in dianion (CL-2) and monoanion (CL-1) ionization states. **(D)** In ideal systems, the changing of ionization state of CL from −1 to −2 causes a minor increase in membrane stiffness and a major reduction in spontaneous curvature. Absence of CL increases membrane stiffness and reduces spontaneous curvature as determined through Martini CG-MD. **(E)** Accounting for headgroup adaptations in complex systems, simulations still show the same trend with dianionic CL and loss of CL showing increases in membrane stiffness and a reduction in spontaneous curvature. **(F)** While increasing lipid saturation has a minimal effect on membrane stiffness in the absence of CL, the increase in the magnitude of spontaneous curvature suggests that the presence of PE can partially, but not completely, compensate for loss of curvature provided by CL. **(G)** Modeling of outer leaflet enrichment of CL in the yeast IMM. Simulated changes in CL concentrations previously reported (Gallet *et al*, 1997) results in membrane softening and increased spontaneous curvature. Two sets of simulations were set up with the estimated compositions of the outer and inner IMM leaflets shown in the pie charts. The simulated outer membrane systems were softer (lower stiffness) and had a larger negative spontaneous curvature. The difference in the outer and inner leaflet curvatures was 0.1±0.04 nm^-1^, which is an estimation for asymmetry-induced *c*_0_.

**Figure EV4:**
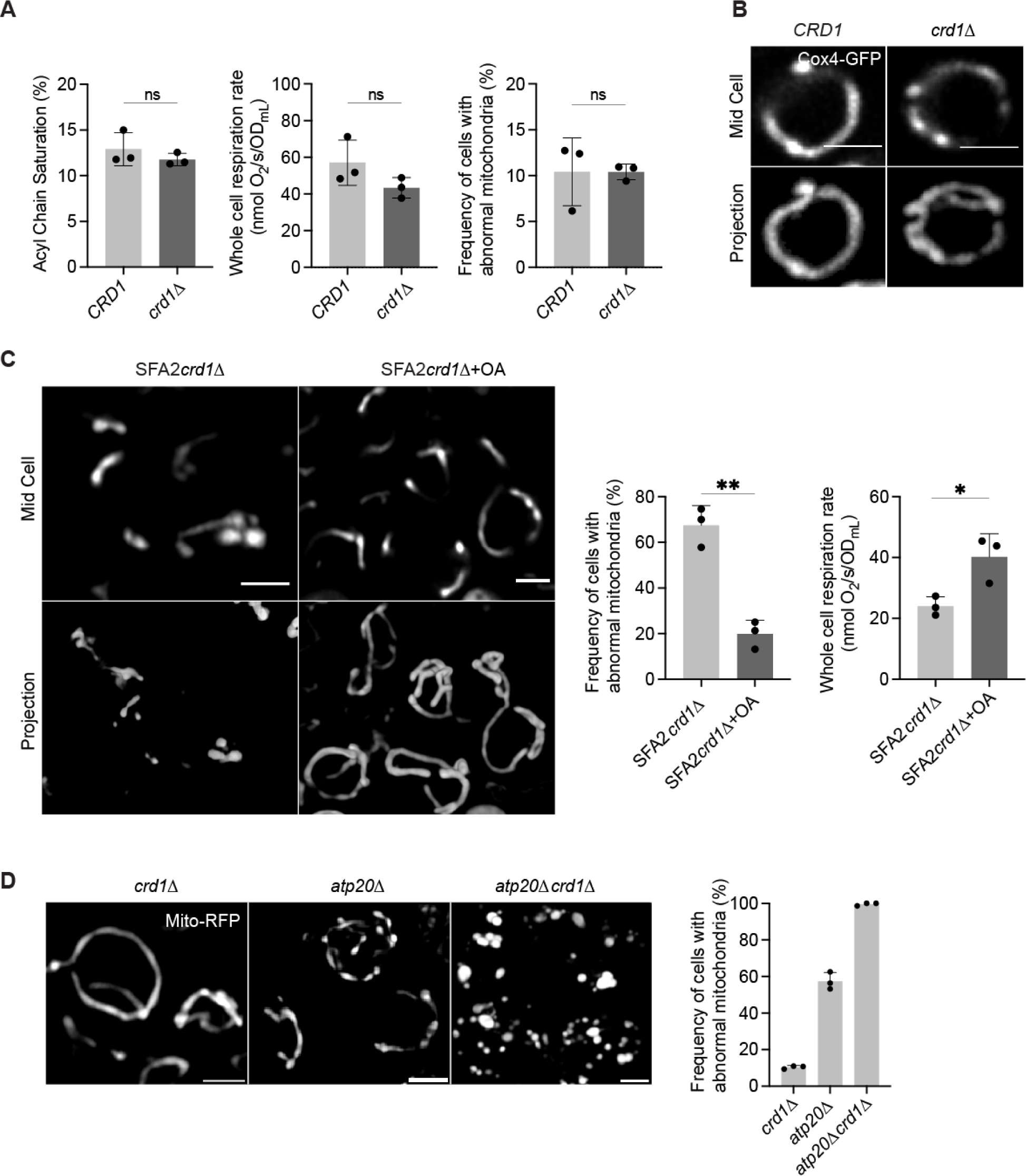
Interactions between cardiolipin synthase, lipid saturation, and ATP synthase dimerization. **(A)** Mitochondrial PL saturation, respiration rate and mitochondrial abnormality measurements of *CRD1* and *crd1*Δ cells. PL saturation was computed from lipidomics analysis on isolated mitochondria. Respiration measurements were performed using Clark electrode, and mitochondrial abnormalities were determined using confocal microscopy with yeast expressing a matrix-localized mts-RFP (N>50 cells quantified per replicate). Measurements were taken from biological replicates (n=3), error bars indicate SD. **(B)** Airyscan confocal micrographs of aerobic wild-type and *crd1*Δ expressing IMM protein Cox4-GFP. Scale bars, 2 μm. Line profile analysis (below) depicts fluorescent intensity across the indicated mitochondria. **(C)** Representative Airyscan confocal micrographs of SFA2*crd1*Δ yeast, grown in the presence and absence of OA, expressing mts-RFP. Scale bars, 2 μm. **p < 0.005, unpaired two-tailed t-test compared against SFA2*crd1*Δ. Respiration rates of SFA2*crd1*Δ cells in the presence and absence of OA were measured in biological replicates (n=3) using a Clark electrode. Error bars indicate SD. *p < 0.05 unpaired two-tailed t-test. **(D)** Loss of CL and ATP synthase dimerization results in complete ablation of mitochondrial morphology and structure as assayed by analysis with mts-RFP. N>50 cells were counted in biological triplicate (n=3) in each condition. Error bars indicate SD. Individual deletion of Crd1p results in normal mitochondrial morphology while half of the cells in *atp20*Δ still retain normal morphology. Scale bars, 2 μm.

**Figure EV5:**
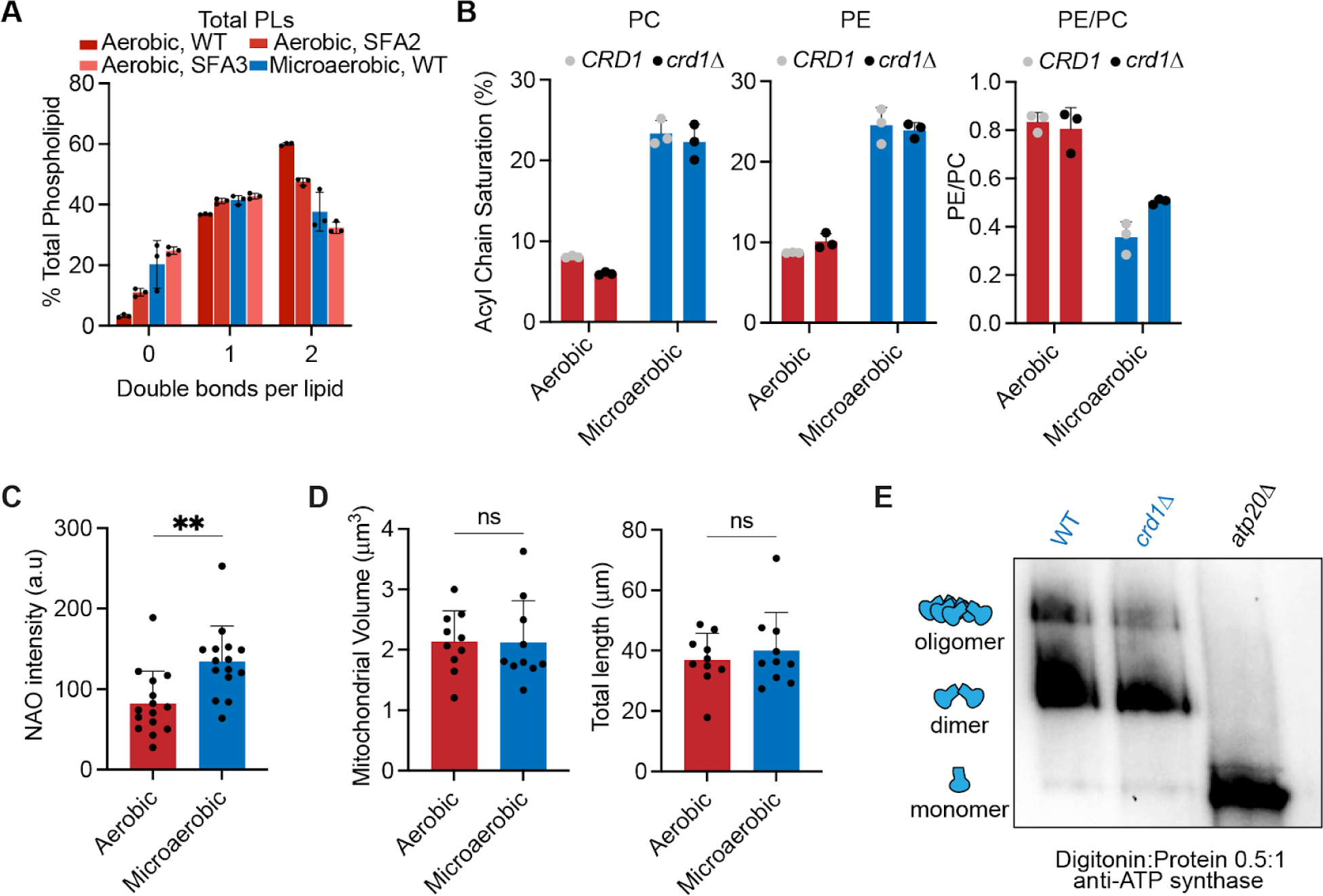
Microaerobic cells exhibit an intermediate level of lipid saturation between SFA2/3 and retain mitochondrial volume and ATP synthase dimers. **(A)** Lipidomic analysis of the double bond distribution of microaerobic yeast (n=3 biological replicates) shows decreased di-unsaturated PL chains and increased saturated PL chains. This is consistent with an intermediary increase in PL saturation between SFA2 and SFA3 levels. Error bars indicate SD. **(B)** The major cellular PL classes, PC and PE, show increased lipid saturation under microaerobic conditions (n=3 biological replicates). Also shown is the decrease in whole cell PE/PC ratio under microaerobic conditions, as observed in SFA strains (Figure 3D). Error bars indicate SD. **(C)** CL content in microaerobic vs. aerobic conditions assayed by staining with 100 nM nonyl acridine orange (NAO). NAO intensities were quantified by line profile analysis from confocal micrographs of N>15 cells in each condition. \*\**p* = 0.0021 unpaired two-tailed t-test against WT. Error bars indicate SD. **(D)** Microaerobic growth conditions do not induce changes to mitochondrial volume or length. Mitograph software was used to measure the voxel volume and total length of mitochondrial tubules from confocal images taken from yeast cells grown in microaerobic vs. aerobic conditions. N=10 cells were quantified from each condition. Error bars indicate SD. **(E)** Microaerobic cells still contain ATP synthase dimers. Digitonin-solubilized crude mitochondria from microaerobic cultures were separated by BN-PAGE and immunoblotted with anti-ATP synthase antibodies. Crude mitochondria from microaerobic *atp20*Δ was used as a monomeric control.

**Figure EV6:**
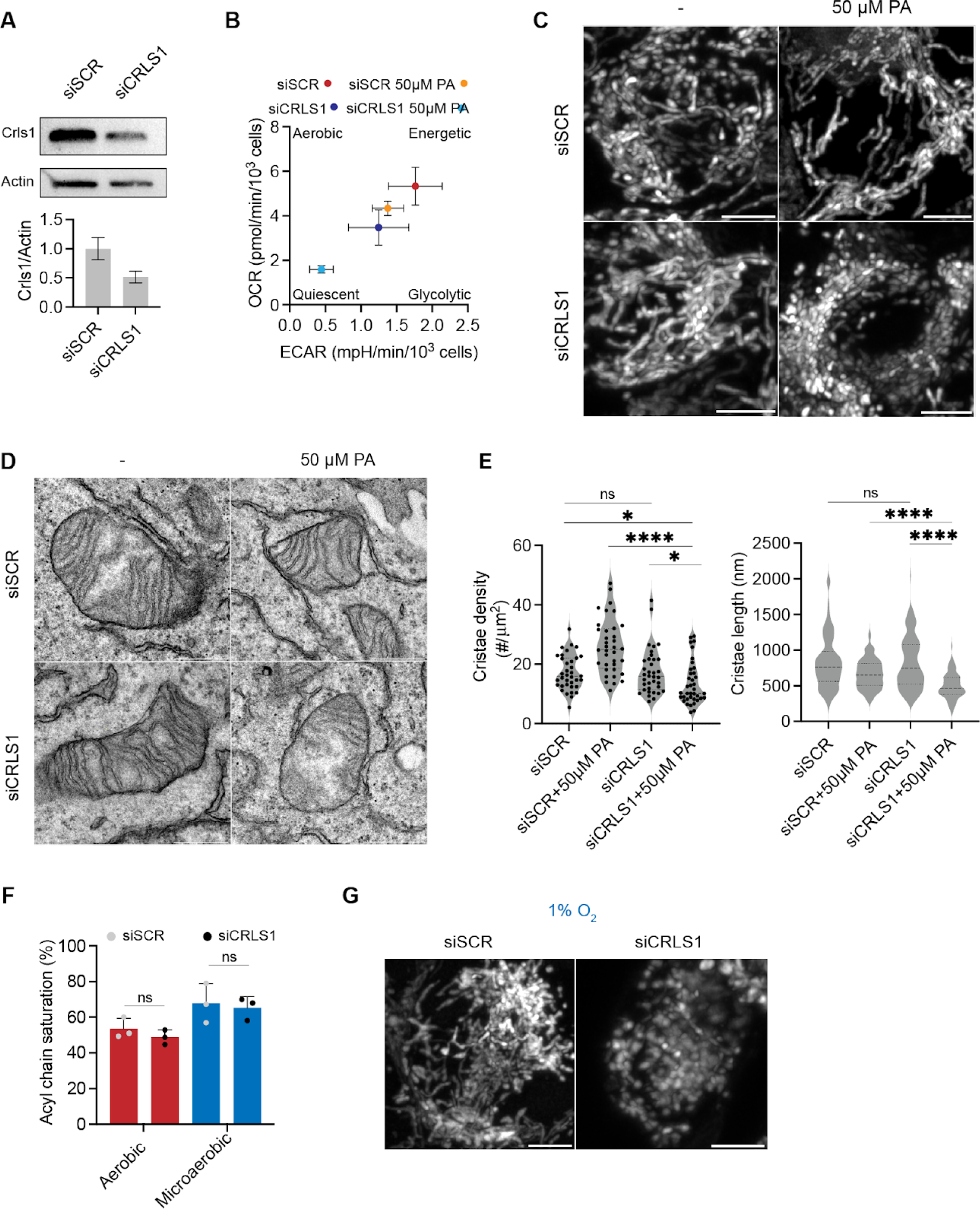
Epistasis between lipid saturation and cardiolipin metabolism in HEK293 cells. **(A)** Knockdown of *CRLS1* by siRNA (siCRLS1) results in a 2-fold decrease in CRLS1 as normalized to β-Actin intensity compared to the scrambled control (siSCR). **(B)** Coordinated increase in saturated fatty acid and depletion of CL result in decreased oxygen consumption rate (OCR) and extracellular acidification rate (ECAR). OCR and ECAR were measured from 10,000 cells in triplicates (n=3) in each condition assayed using Seahorse respirometry. Cells treated with both mild concentrations (50 μM) of PA and siCRLS1 showed a quiescent metabolic phenotype. **(C)** Knockdown of *CRLS1* results in loss of tubular mitochondrial morphology upon treatment with 50 μM PA. Representative images of single cells transfected with scrambled (siSCR) or *CRLS1* (siCRLS1) siRNAs and subsequently treated with BSA or BSA complexed with 50μM PA. Cells were stained with Mitotracker DeepRed. Images show individual cells. Scale bars, 5 μm. **(D)** *CRLS1* knockdown cells treated with 50 μM PA are bereft of abundant cristae sheets observed in control cells or those treated with only siCRLS1 or 50μM PA. Representative TEMs of individual mitochondria are shown from each condition. Scale bars, 500 nm. **(E)** *CRLS1* knockdown cells treated with 50 μM PA show decreased cristae density and cristae length. Cristae density was quantified as the number of cristae per mitochondrial area and assessed from N=35 mitochondria. ****p < 0.0001 unpaired t-test of siCRLS1 with 50 μM PA against siSCR with 50μM PA, *p<0.025 unpaired t-test of siCRLS1 with 50μM PA against siSCR and siCRLS1. Cristae length was measured in N=100 cristae in each condition. ****p < 0.0001 unpaired t-test of siCRLS1 with 50μM PA against siSCR with 50μM PA and siCRLS1. Error bars indicate SD. **(F)** Knockdown of *CRLS1* does not change the acyl chain saturation in aerobic or microaerobically grown cells, but the latter feature higher lipid saturation due to reduced oxygenation. Error bars indicate SD (n=3). Error bars indicate SD. **(G)** Under 1% oxygen growth conditions, knockdown of *CRLS1* results in loss of tubular mitochondrial morphology. Cells were incubated in microaerobic chambers for 72 hours or normoxic chambers for 48 hours prior to staining with Mitotracker DeepRed. Images show individual cells. Scale bars, 5 μm.

**Appendix Figure S1:**
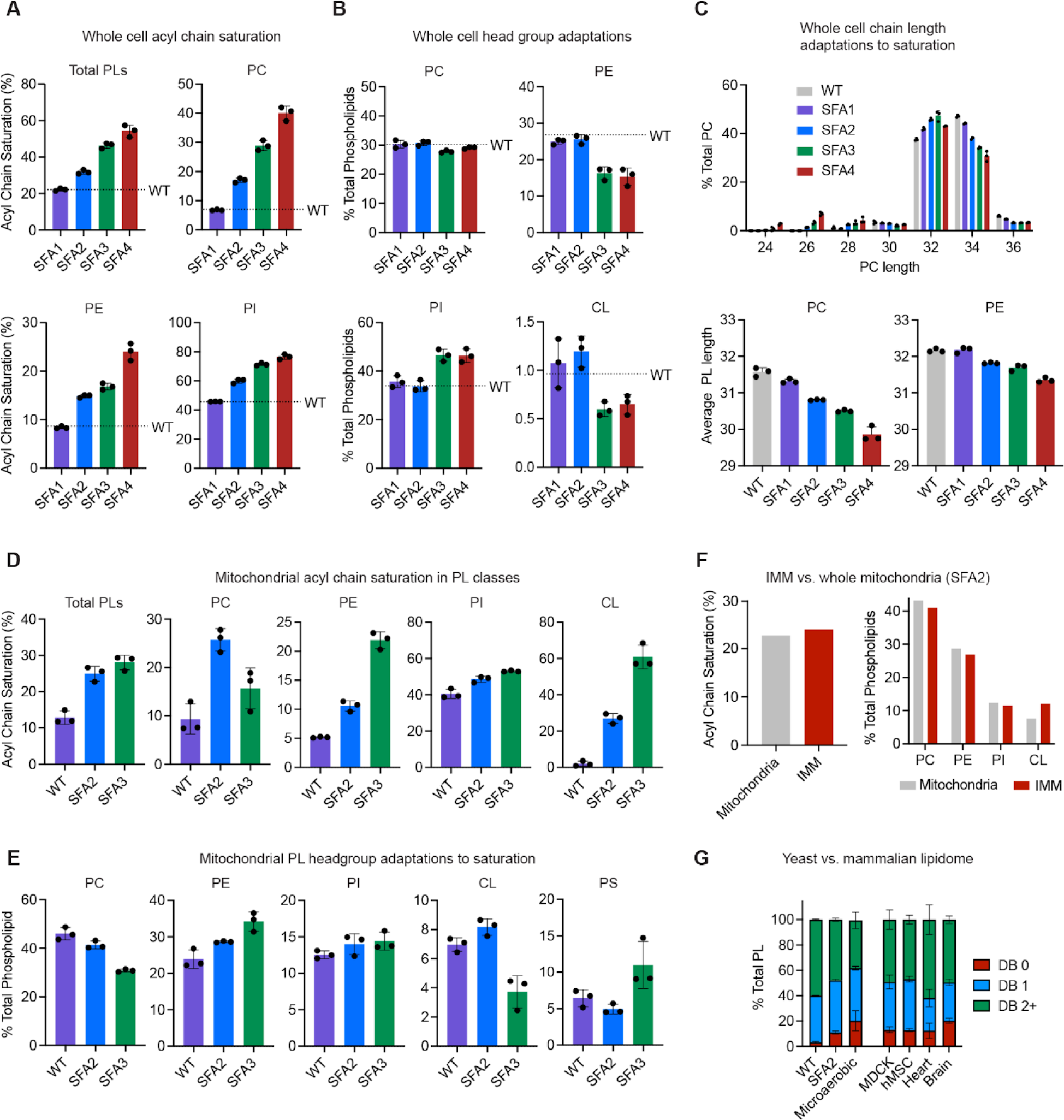
Changes to whole cell and mitochondrial lipid profile upon modulation of Ole1p expression. **(A)** Decreasing Ole1p expression increases the acyl chain saturation of the total yeast PL pool, and major PL classes as assayed by lipidomics, error bars indicate SD, n=3 independent cultures. WT levels are shown as dotted lines. **(B)** Potentially compensatory changes to the whole cell lipidome in response to increasing saturation. Shown are the abundance of major yeast PLs in SFA strains, n=3 independent cultures. WT levels shown as dotted lines. As saturation increases, PE and CL decrease, while PI increases. Error bars indicate SD. **(C)** Increasing lipid saturation results in shortening of PC and PE acyl chains in the whole cell. Shown are the sum of the lengths for the *sn*-1 and *sn*-2 chains. **(D)** Acyl chain saturation in isolated mitochondria from SFA strains and WT as determined by intact lipid analysis for the total PL pool, n=3 independent cultures. Error bars indicate SD. **(E)** PL headgroup adaptations in isolated mitochondria from SFA strains and wild-type in major PL classes as determined by lipidomic analysis, n=3 independent cultures. In the mitochondria, PC decreases and PE increases as saturation increases. Error bars indicate SD. **(F)** The IMM and whole mitochondrial lipidome display similar levels of saturation and PE/PC, but differ in abundance of CL (higher in the IMM). **(G)** Mammalian lipidomes contain similar lipid saturation profiles to yeast SFA2 and micro-aerobically grown cells, where CL is essential. Shown is the double bond profile of all PLs as determined through lipidomics from *S. cerevisiae* (this study) in comparison with mammalian cell lines (MDCK-CM and hMSC-CM) and isolated tissues previously analyzed using the same lipidomics platform(Symons *et al*, 2021). WT yeast grown under vigorous aeration have a low number of saturated and monounsaturated PLs, while WT cells grown under physiologically-relevant oxygen concentrations (microaerobic) or engineered strains (SFA2) show profiles more similar to mammalian cells. In the latter systems, CL is an essential component for proper mitochondrial biogenesis. Error bars indicate SD.

**Appendix Figure S2:**
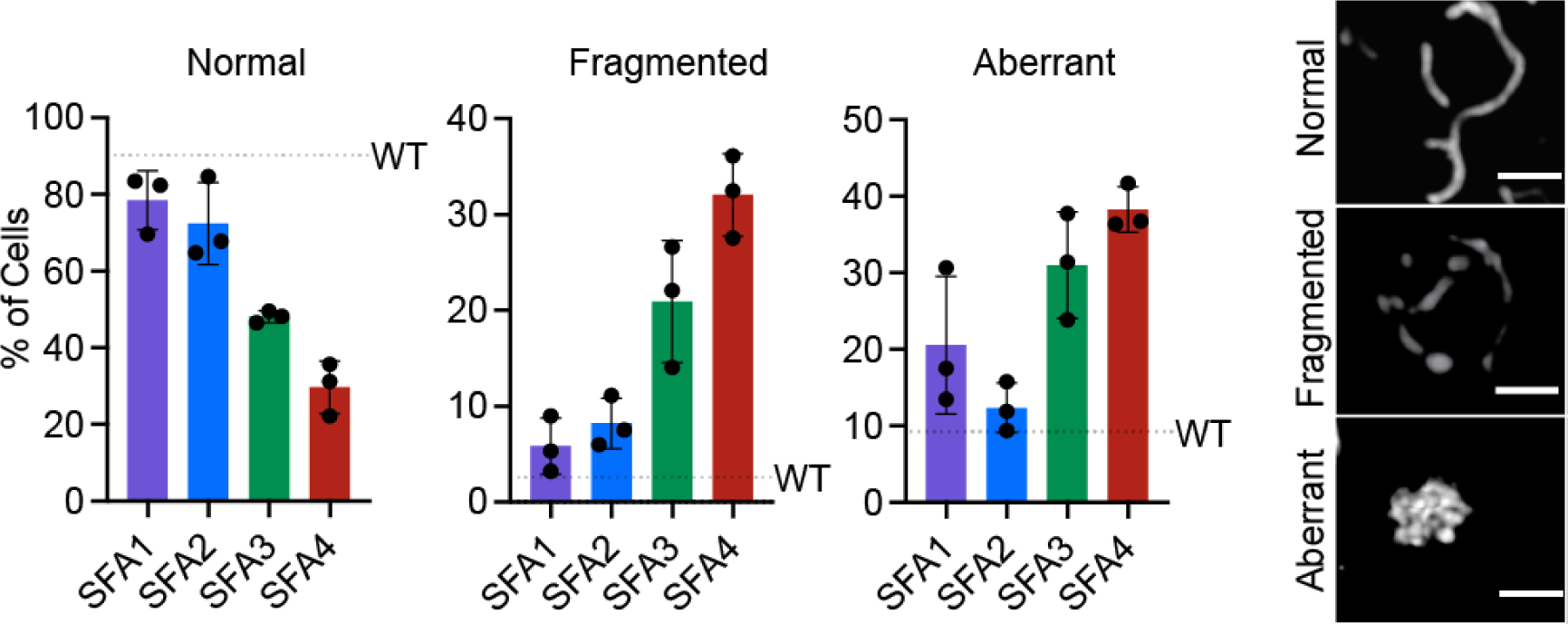
Pipeline for quantifying changes in mitochondrial morphology observed in SFA strains. Imaging of matrix-localized RFP was used to quantify the frequency of mitochondrial abnormality as a function of the amounts of normal, fragmented and aberrant mitochondria. Normal mitochondria contain tubulations throughout the whole yeast cell, while fragmented mitochondria retain an overall mitochondrial structure but have lost the interconnected tubular network associated with normal mitochondria. Aberrant mitochondria are characterized by punctate aggregations of mitochondria in the center of the yeast cell. Cells were imaged in n=3 independent cultures. Error bars indicate SD. Scale bars, 2 μm.

**Appendix Figure S3:**
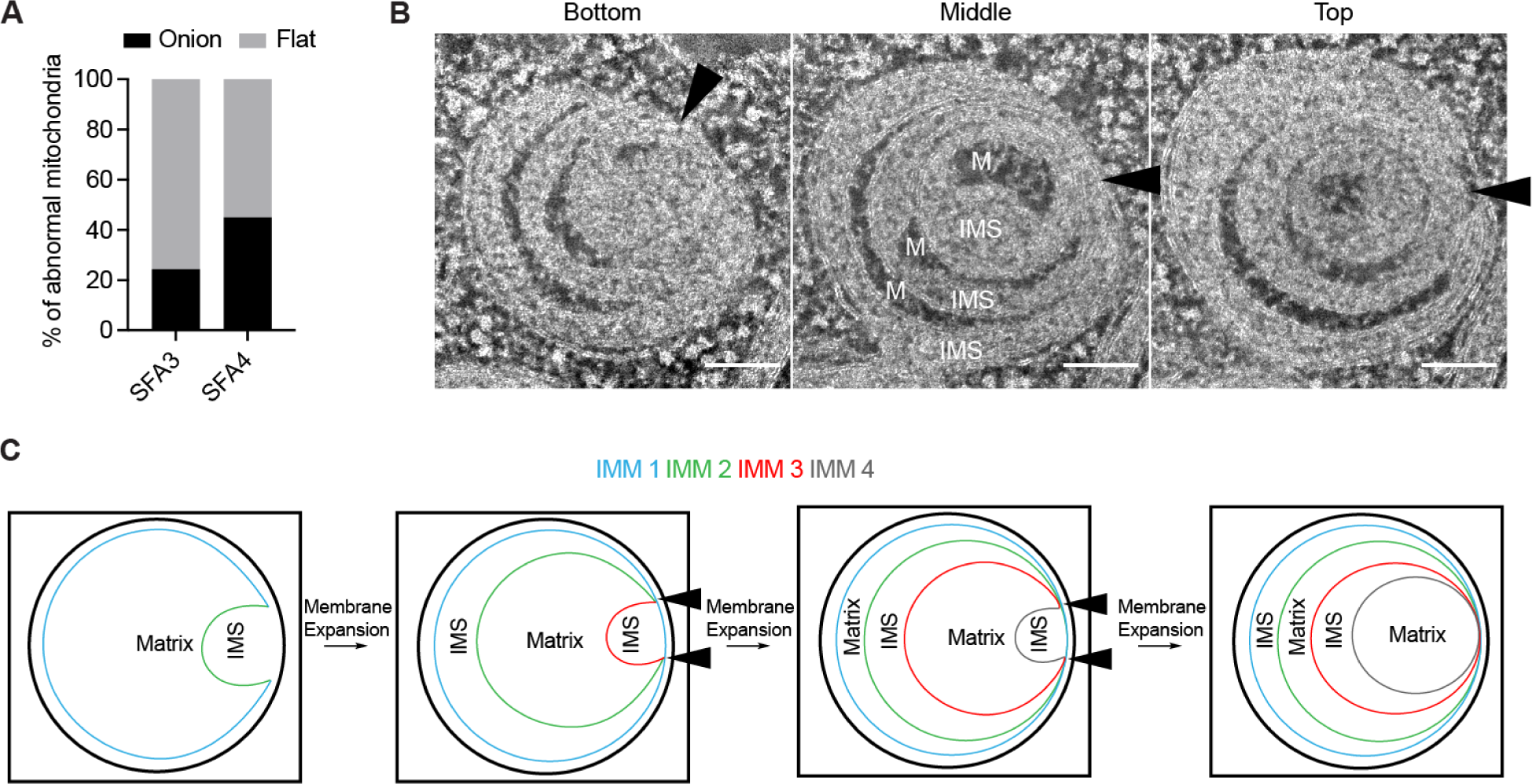
Multi-tilt electron tomography reveals a mechanism for onion-formation in aberrant mitochondria. **(A)** Quantitative analysis of thin-section TEM micrographs reveals the abnormal mitochondria in SFA3 are predominantly flat, while in SFA4 there is an even distribution of onion and flat abnormal mitochondria. At least N=40 mitochondria were quantified from each strain. **(B)** Multi-tilt tomogram slices of HPFS SFA4 yeast cells at three z-positions. ‘M’ indicates matrix regions (dark), while ‘IMS’ indicates intermembrane space regions (light). Shading indicates alternating matrix and IMS regions, as previously observed (Paumard *et al*, 2002). Black triangles indicate observed regions of contact points between IMM layers, suggesting a continuous IMM. Scale bars, 100 nm. **(C)** Schematic depiction for one model of how onion-like morphology could be formed by a continuous IMM undergoing subsequent buddings during its biogenesis due to membrane expansion. Black triangles indicate regions of contact points between multiple apparent IMM layers that are continuous.

**Appendix Figure S4:**
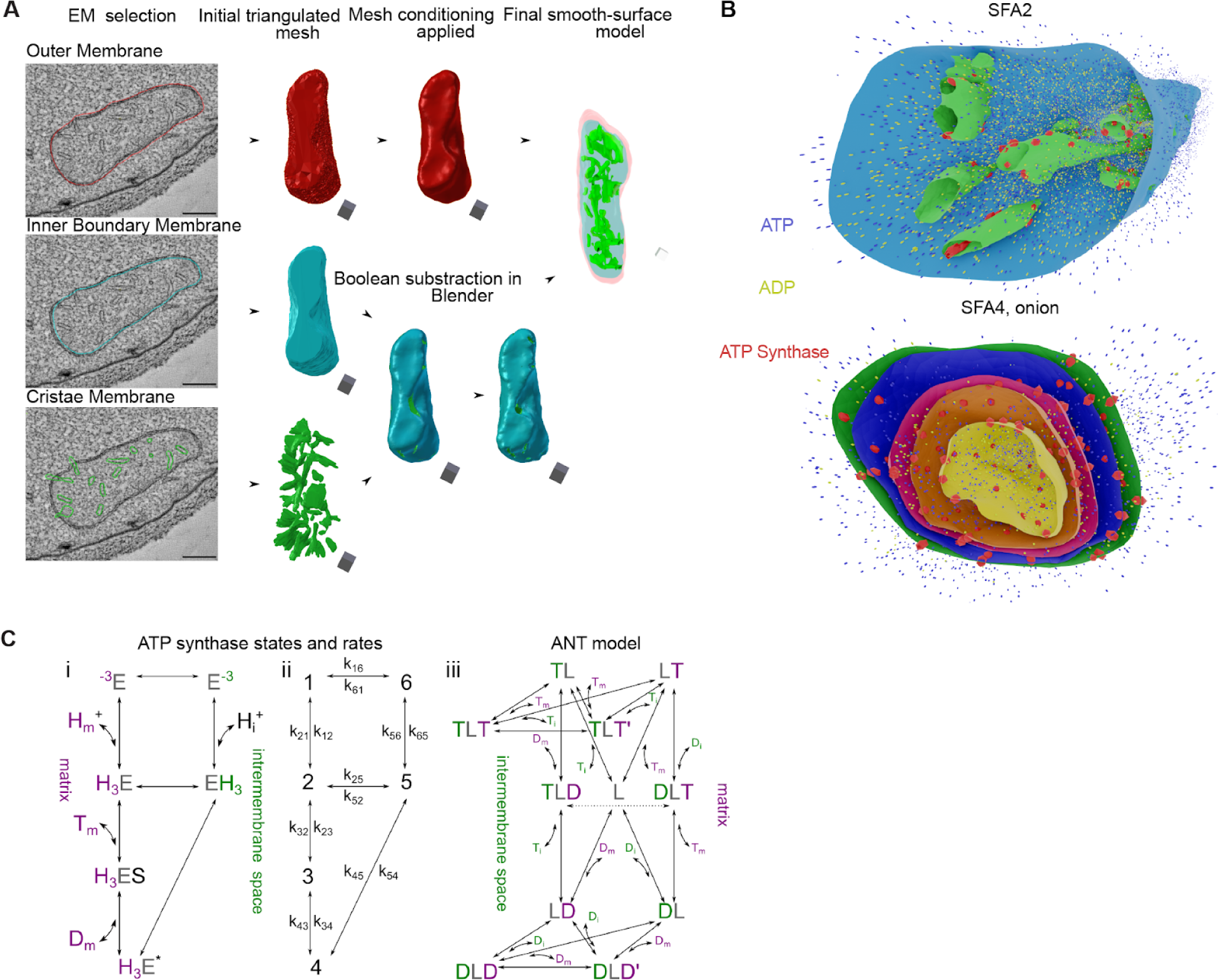
Pipeline for mesh generation from multi-tilt tomography for ATP synthesis simulations. **(A)** Example of Blender-based 3D Mesh generation pipeline from EM tomograms. **(B)** Snapshots of ATP generation simulations as displayed in Movie S4 and S5. SFA2 shows localizations of ATP synthases to regions of high curvature in CMs, while in the SFA4 onion ATP synthases are distributed evenly on each layer of IMM based on previous cryo electron tomography reconstructions. **(C) (i-iii)** Schematic representations of the kinetic states and modeled rates of ATP synthase and ANT used to construct the metabolic model. Further details of the model can be found in the appendix modeling procedures.

**Appendix Figure S5:**
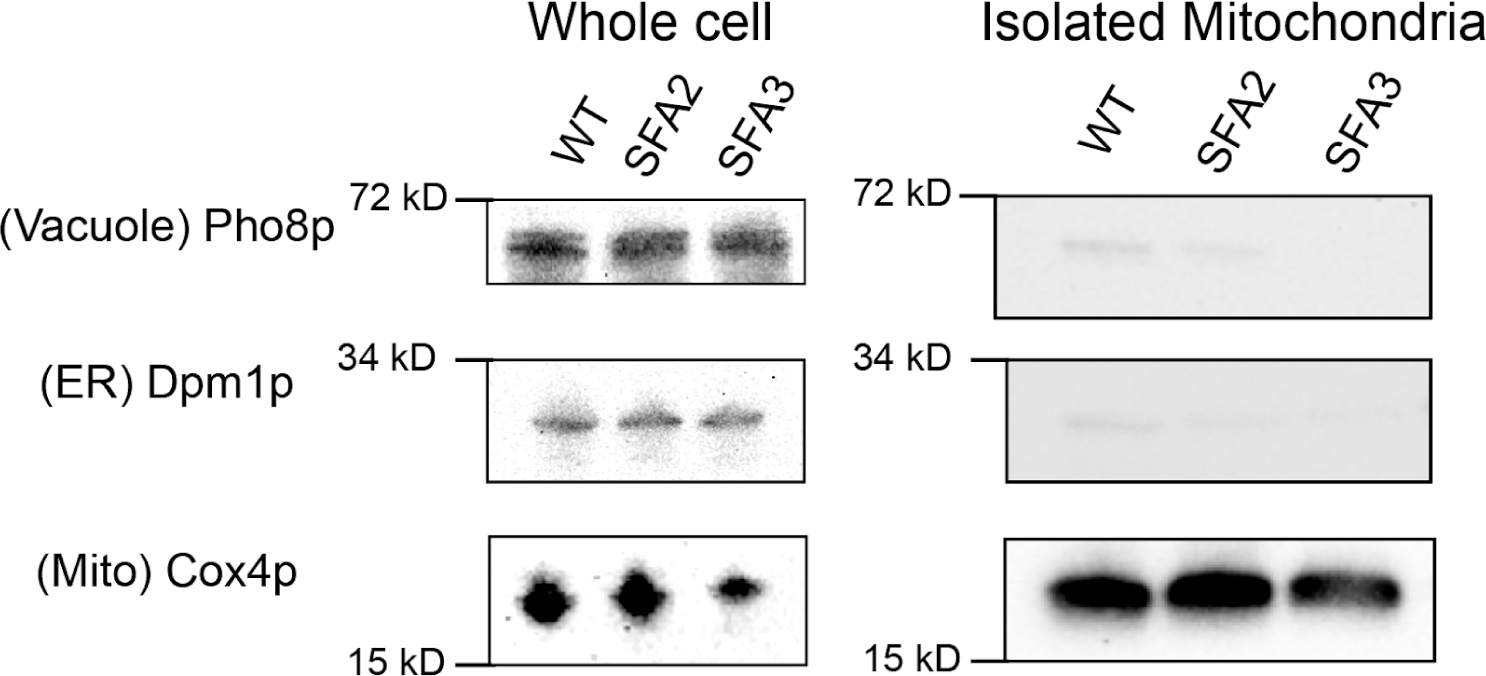
Isolated mitochondrial fractions are bereft of contamination from other organelles. Whole cell lysates were first grown to the stationary phase in YPEG. 2.5 OD units of cells were taken and subjected to protein extraction and SDS-PAGE as previously described (Kushnirov, 2000). Gels were then transferred and western blotted before decoration with antibodies against the vacuole, ER and mitochondria. For isolated mitochondria, 10μg of protein was loaded and subjected to western blotting against organelle antibodies. For each antibody, the following dilutions were used: 1:1000 for Cox4p and Dpm1p and 1:250 for Pho8p.

**Appendix Figure S6:**
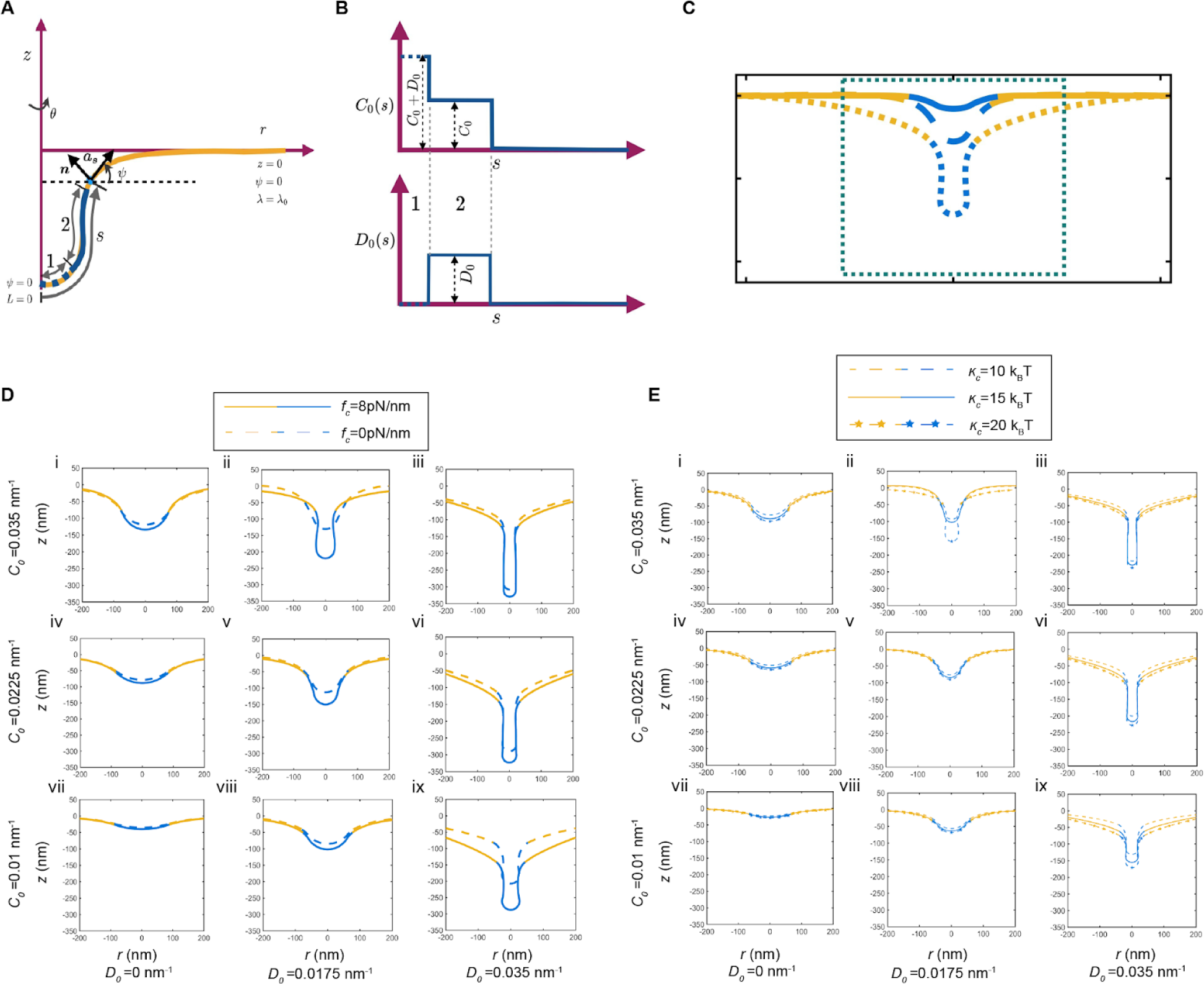
Continuum modeling details and comparison of tubular morphologies with and without an applied collar force. **(A)** Schematic showing the axisymmetric membrane configuration along with the boundary conditions. The yellow regions depict the bare lipid bilayer, and the blue regions depict the regions where different spontaneous curvatures are prescribed. **(B)** Prescription of isotropic and anisotropic spontaneous curvature along the arc length in the simulations. **(C)** The simulation domain is large to avoid boundary effects but the zoomed in portion in the dashed box is shown to demonstrate the shapes of the membrane. **(D)** Comparison of the tubular shapes with (solid lines) and without (dashed line) the collar force at the base of the cristae. All parameters are the same as those in Figure 5B. Presence of a collar force promoted cristae like structures for the same values of imposed curvatures. This is particularly apparent in panel ii. **(E)** Shapes of the membrane for the same values of coat area, collar force, and variations in the isotropic and anisotropic spontaneous curvature as panel D for different values of bending modulus. We observe that the shapes of the tubular cristae are not sensitive to changes in bending modulus for low curvature values but differences in membrane curvature can be seen for high values of isotropic and anisotropic spontaneous curvatures.

### Supplementary Tables

**Table S1A:** Bending moduli (*к_c_*) values extracted from X-ray scattering analysis or micropipette aspiration analysis of PC membranes as a function of acyl chain saturation.

X-ray scattering
| Lipid | Acyl Chains | $\kappa_c$ ($10^{-20}$ J) | Source |
| --- | --- | --- | --- |
| POPC | 16:0 18:1 | 8.5 | (Kucerka *et al*, 2005) |
| DOPC | 18:1 18:1 | 8.5±0.6 | (Jablin *et al*, 2014) |

Micropipette aspiration
| Lipid | Acyl Chains | $\kappa_c$ ($10^{-19}$ J) | Source |
| --- | --- | --- | --- |
| SOPC | 18:0 18:1 | 0.90±0.06 | (Rawicz *et al*, 2000) |
| DOPC | 18:1 18:1 | 0.85±0.10 | (Rawicz *et al*, 2000) |

**Table S1B:**
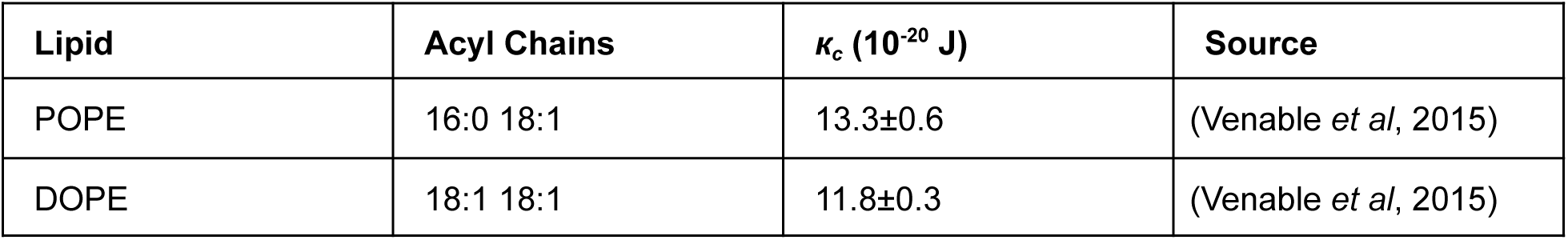
Bending moduli (*к_c_*) values extracted from MD simulations of PE as a function of acyl chain saturation.

**Table S1C:** Spontaneous curvature (*c_0_*) values extracted from SAXS analysis of PC and PE lipids as a function of acyl chain saturation.

PC
| Lipid | Acyl Chains | Matrix | $c_0$ ($\text{nm}^{-1}$) | Source |
| --- | --- | --- | --- | --- |
| DPPC | 16:0 16:0 | DOPE | 0.05±0.05 | (Kaltenegger *et al*, 2021) |
| POPC | 16:0 18:1 | DOPE | 0.01±0.04 | (Kaltenegger *et al*, 2021) |
| DOPC | 18:1 18:1 | DOPE | -0.04±0.04 | (Kaltenegger *et al*, 2021) |

PE
| Lipid | Acyl Chains | Matrix | $c_0$ ($\text{nm}^{-1}$) | Source |
| --- | --- | --- | --- | --- |
| POPE | 16:0 18:1 | - | -0.317±0.007 | (Frewein *et al*, 2019) |
| DiPoPE | 16:1 16:1 | - | -0.382±0.009 | (Frewein *et al*, 2019) |
| DOPE | 18:1 18:1 | - | -0.409±0.010 | (Frewein *et al*, 2019) |

**Table S2:**
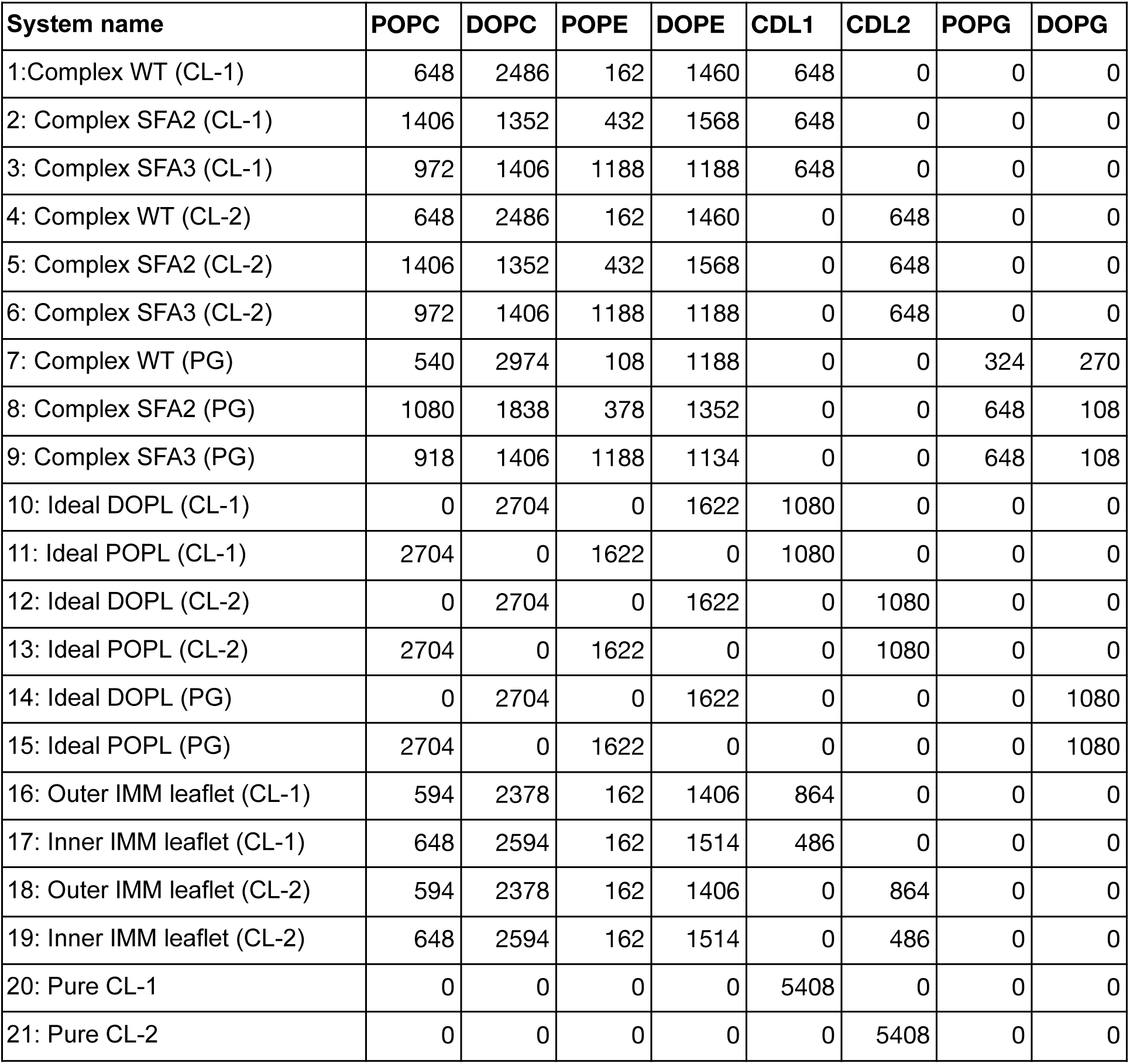
List of membrane compositions simulated by CG-MD; the number of lipids for each type are shown.

| System name | POPC | DOPC | POPE | DOPE | CDL1 | CDL2 | POPG | DOPG |
| --- | --- | --- | --- | --- | --- | --- | --- | --- |
| 1: Complex WT (CL-1) | 648 | 2486 | 162 | 1460 | 648 | 0 | 0 | 0 |
| 2: Complex SFA2 (CL-1) | 1406 | 1352 | 432 | 1568 | 648 | 0 | 0 | 0 |
| 3: Complex SFA3 (CL-1) | 972 | 1406 | 1188 | 1188 | 648 | 0 | 0 | 0 |
| 4: Complex WT (CL-2) | 648 | 2486 | 162 | 1460 | 0 | 648 | 0 | 0 |
| 5: Complex SFA2 (CL-2) | 1406 | 1352 | 432 | 1568 | 0 | 648 | 0 | 0 |
| 6: Complex SFA3 (CL-2) | 972 | 1406 | 1188 | 1188 | 0 | 648 | 0 | 0 |
| 7: Complex WT (PG) | 540 | 2974 | 108 | 1188 | 0 | 0 | 324 | 270 |
| 8: Complex SFA2 (PG) | 1080 | 1838 | 378 | 1352 | 0 | 0 | 648 | 108 |
| 9: Complex SFA3 (PG) | 918 | 1406 | 1188 | 1134 | 0 | 0 | 648 | 108 |
| 10: Ideal DOPL (CL-1) | 0 | 2704 | 0 | 1622 | 1080 | 0 | 0 | 0 |
| 11: Ideal POPL (CL-1) | 2704 | 0 | 1622 | 0 | 1080 | 0 | 0 | 0 |
| 12: Ideal DOPL (CL-2) | 0 | 2704 | 0 | 1622 | 0 | 1080 | 0 | 0 |
| 13: Ideal POPL (CL-2) | 2704 | 0 | 1622 | 0 | 0 | 1080 | 0 | 0 |
| 14: Ideal DOPL (PG) | 0 | 2704 | 0 | 1622 | 0 | 0 | 0 | 1080 |
| 15: Ideal POPL (PG) | 2704 | 0 | 1622 | 0 | 0 | 0 | 0 | 1080 |
| 16: Outer IMM leaflet (CL-1) | 594 | 2378 | 162 | 1406 | 864 | 0 | 0 | 0 |
| 17: Inner IMM leaflet (CL-1) | 648 | 2594 | 162 | 1514 | 486 | 0 | 0 | 0 |
| 18: Outer IMM leaflet (CL-2) | 594 | 2378 | 162 | 1406 | 0 | 864 | 0 | 0 |
| 19: Inner IMM leaflet (CL-2) | 648 | 2594 | 162 | 1514 | 0 | 486 | 0 | 0 |
| 20: Pure CL-1 | 0 | 0 | 0 | 0 | 5408 | 0 | 0 | 0 |
| 21: Pure CL-2 | 0 | 0 | 0 | 0 | 0 | 5408 | 0 | 0 |

**Table S3:**
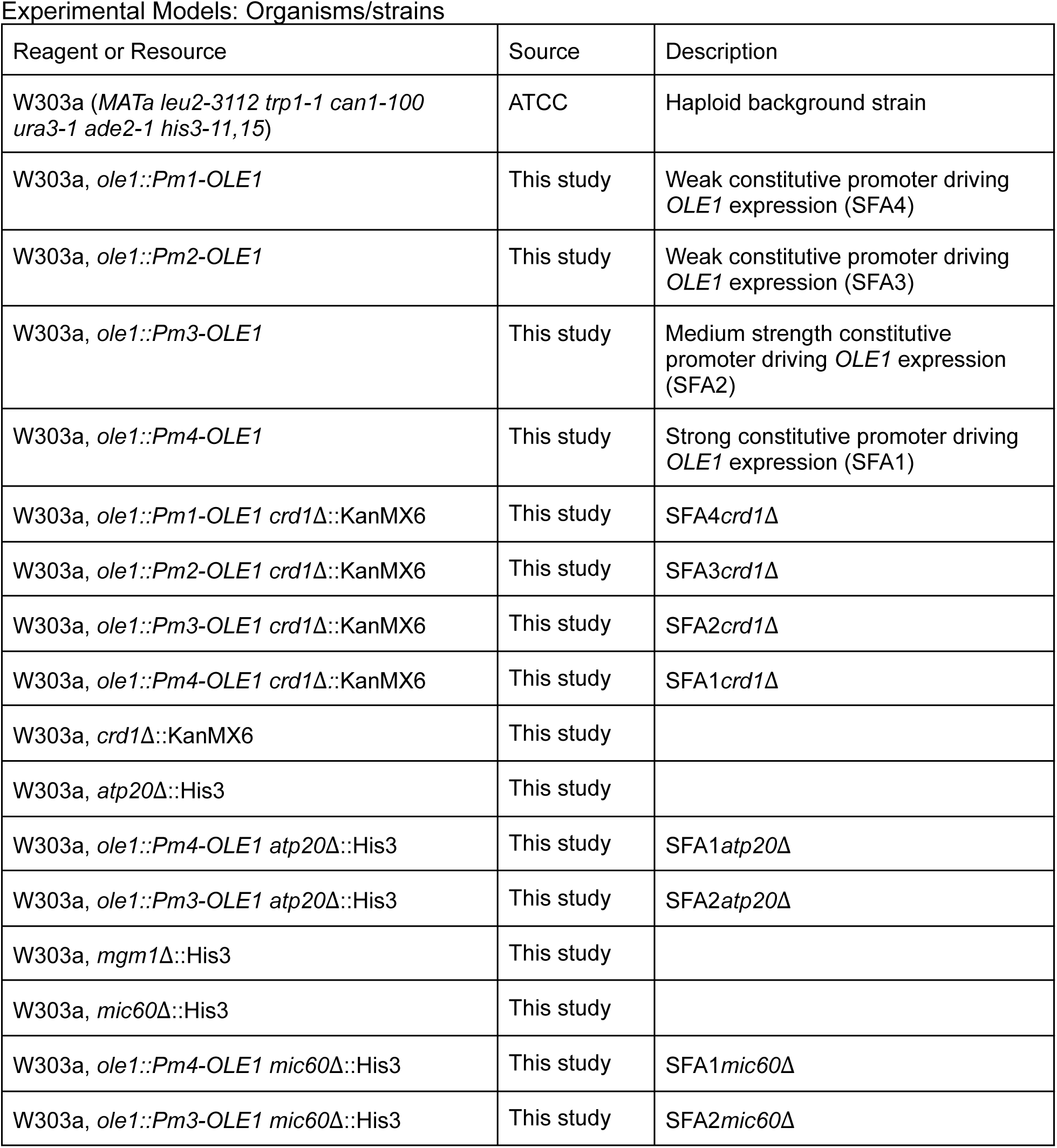
Strains used in this study Experimental Models: Organisms/strains.

**Table S4:**
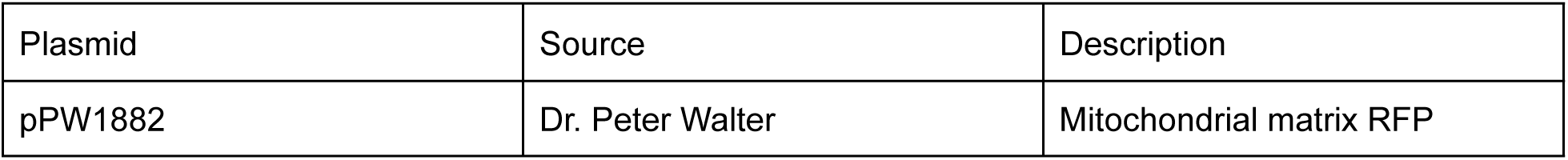
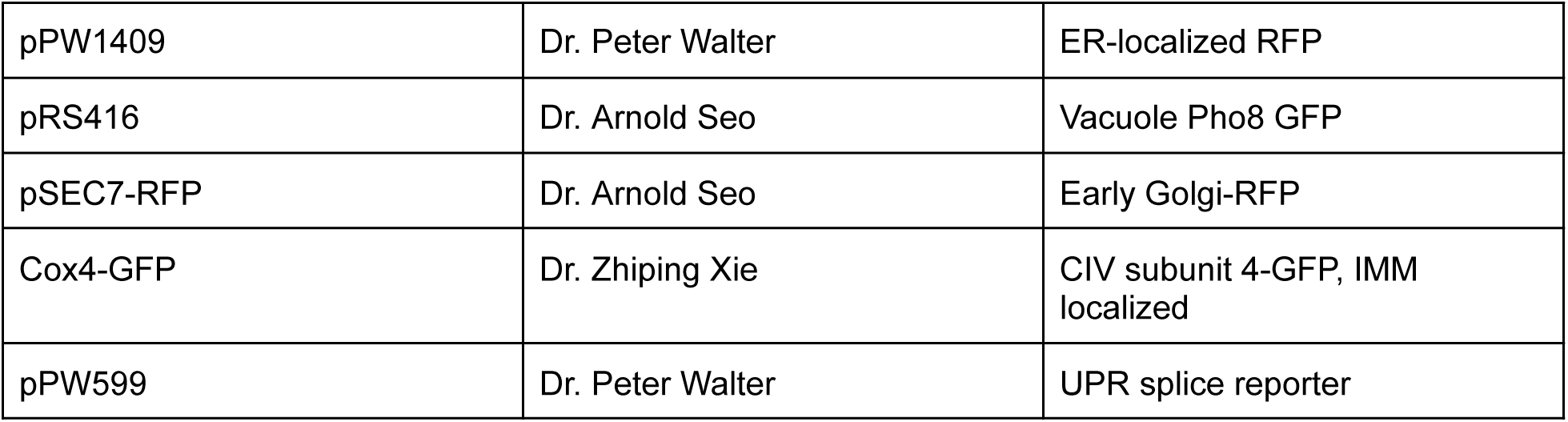
Plasmids used in this study.

**Table S5:** Antibodies used in this study.

| Reagent or Resource | Source | Description |
| --- | --- | --- |
| Anti-ATP synthase subunit $\beta$ antibody | Dr. Alexander Tzagoloff | Goat anti-rabbit, polyclonal (Rak & Tzagoloff, 2009) |
| Anti-Mic60p | Dr. Andreas Reichart | Goat anti-rabbit (Rabl *et al*, 2009) |
| Anti-Mgm1p | Dr. Andreas Reichart | Goat anti-rabbit (Rabl *et al*, 2009) |
| Anti-Cox4p | Abcam | Cat: 110272 |
| Anti-Dpm1p | Abcam | Cat: 113686 |
| Anti-Pho8 | Abcam | Cat: 113688 |
| Anti-Cox1 | Abcam | Cat: 11D8B7 |
| Anti-CRLS1 | Proteintech | Cat: 14845-1-AP |
| Anti-Actin | Invitrogen | Cat: PA5-78715 |

### Multimedia file captions

**Movie S1:** Animation depicting the mean curvature map within the IMM of a cristae-containing SFA2 mitochondrion.

**Movie S2:** Animation depicting the mean curvature map within the flat IMM of a SFA4 mitochondrion.

**Movie S3:** Animation depicting the mean curvature map within the IMM of an onion-like SFA4 mitochondrion.

**Movie S4:** Animation depicting the simulated distribution of ATP synthases (large red spheres), ANTs, ATP (small blue spheres) and ADP (small yellow spheres) within the IMM of a cristae-containing SFA2 mitochondrion.

**Movie S5:** Animation depicting the simulated distribution of ATP synthases (large red spheres), ANTs, ATP (small blue spheres) and ADP (small yellow spheres) within the IMMs of an onion-like SFA4 mitochondrion.

**Movie S6:** Animation showing the change in the shape of membrane for bending modulus 15 k_B_T, with collar force 8 pN/nm, and D_0_ fixed at 0.0175 nm^-1^. The movie shows the shape as C_0_ increases.

**Movie S7:** Animation showing the change in the shape of membrane for bending modulus 15 k_B_T, with collar force 8 pN/nm, and C_0_ fixed at 0.0225 nm^-1^. The movie shows the shape as D_0_ increases.

### Modeling Procedures

#### ATP production modeling

The computational for ATP generation in mitochondria is based on previous modeling efforts (Garcia *et al*, 2019, 2022; Magnus & Keizer, 1997; Bertram *et al*, 2006; Saa & Siqueira, 2013). We solve the reactions (detailed below) using MCell (Kerr *et al*, 2008) to accurately capture the stochastic nature of the events underlying ATP production in the small volumes of the mitochondria. The model has a total of 19 equations and 41 parameters and the thermodynamic details are given in (Garcia *et al*, 2022). We briefly describe the main components of the model below.

##### ATP synthase

The ATP synthase model is composed of ATP synthase (represented as E) that can be in six states (Figure S3F), representing different protein configurations. Each state corresponds with a number from 1 to 6, and each transition has associated a rate constant k_ij_ (transition from the state i → j). In some cases k_ij_ depends on the membrane potential, proton concentration, or phosphate concentration. The list of reactions and model parameters are given below, reproduced from(Garcia *et al*, 2022). The model was adapted from the work of Pietrobon and Caplan(Pietrobon & Caplan, 1985).

ATP synthase is modeled as a membrane protein that can transport protons (H^+^) from and to the matrix and synthesize ATP. The translocation of 3 H^+^ is coupled to the phosphorylation of one ADP into ATP, approximating the stoichiometry of the yeast ATP synthase with a c_10_ ring (10 H^+^/ 3 ATP). The free enzyme with its negative charged cavity facing the IMS is represented by E^-3^. Three protons can bind, generating the transition to state EH_3_. The protons can be translocated to the matrix through the reaction (EH_3_ → H_3_E) or EH_3_ → H_3_E*. A transition to state H_3_ES can follow binding one ADP molecule from the matrix (represented as D_m_) under constant phosphate (P_i_) concentration, which is kept at 20 mM. This is followed by the production of one molecule of ATP (T_m_) through the reaction H_3_ES -> H_3_E + T_m_. Finally, in the transition H^3^E → ^-3^E + 3H^+^ three protons are unbound in the matrix. The negative charged cavity of the enzyme can also transition from facing the matrix (state ^-3^E) to facing the IMS (state E^-3^).

Transition 6 → 5 accounts for the binding of 3 protons from the IMS to the free enzyme (state 6, E^−3^), two transitions can occur from here: transition 5 → 4 represents the transport of the protons to the matrix or transition 5 → 2 that represents the transport of the protons to the matrix without producing ATP. In state 4, ADP can bind to the enzyme (transition 4 → 3) and subsequently ATP can be synthesized (transition 3 → 2). This is followed by the unbinding of the protons in the matrix (transition 2 → 1), arriving at state 1.

List of Reactions for the ATP synthase model: (1) ^−3^E + 3H^+^_m_ ↔ H_3_E, k_12_, k_21_ (2) H_3_E + T_m_ ↔ H_3_ES, k_23_, k_32_ (3) H_3_ES ↔ H_3_E^∗^ + D_m_, k_34,_ k_43_ (4) H_3_E^∗^ ↔ EH_3_, k_45_, k_54_ (5) EH_3_ ↔ E^−3^ + H^+^_i_, k_56_, k_65_ (6)E−3 ↔−3E, k61, k16

Parameter values for the ATP synthase model: k_43_ = 2×10^6^ M^−1^ s^−1^, k_34_ = 100 s^−1^, k_12_ = 25 s^−1^, k_21_ = 40 s^−1^, k_65_ = 3969 s^−1^, k_56_ = 1000 s^−1^, k_61_ = 33989 s^−1^, k_16_ = 146516 s^−1^, k_54_ = 1×10^2^ s^−1^, k_45_ = 100 s^−1^, k_25_ = 5.85×10^−30^ s^−1^,k_52_ = 1×10^−20^ s^−1^, k_32_ = 5×10^3^ s^−1^, k_23_ = 5×10^6^ M^−1^ s^−1^

##### Modeled distribution of ATP synthases

The density of ATP synthases has been estimated at 3070 ATP synthases per um^2^ in areas of high membrane curvature (Acehan *et al*, 2011), this is consistent with ATP synthase densities estimated from yeast (Davies *et al*, 2012). For each reconstruction, we calculated the surface area formed by vertices with first principal curvature higher than 70 μm^-1^, and with this the number of ATP synthases was estimated for each organelle. For instance, the surface area of high curvature for the reconstruction of an SFA2 mitochondria is 0.144 μm^2^, which leads to an estimation of 433 units of ATP synthases in this reconstruction. To perform the spatial simulations, ATP synthases were distributed randomly in the regions of high curvature. For each mitochondrion, the total number of ATP synthases was kept the same.

##### ATP/ADP translocator (ANT) model

The model for the ATP/ADP translocator (ANT) is composed of 11 states and 22 chemical reactions, listed below. The kinetic diagram is presented in Figure S3F. The free protein is represented with the letter L in the diagram; it can bind ADP (D) or ATP (T) molecules from the matrix side (on the right) or IMS side (on the left), forming a triple molecular state. State TLD for instance represents a state with one ATP bound from the IMS and one ADP from the matrix side. The reaction that transports ATP from the matrix to the IMS is DLT → TLD, the rate constant for this reaction is k_p_, the reverse reaction imports ATP to the matrix and exports ADP to the IMS, with rate constant k_cp_. Futile translocations can also occur translocating one molecule of ATP by another ATP (TLT → TLT’). TLT and TLT’ represent the same state, but they are differentiated to measure the rate of these translocations.

List of Reactions for the ANT model: (1) T_m_ + L ↔ LT, k^+^_Tm_, k^-^_Tm_ (2) D_m_ + L ↔ LD, k^+^_Dm_, k^-^_Dm_ (3) T_i_ + L ↔ TL,, k^+^_Ti_, k^-^_Ti_ (4) D_i_ + L ↔ DL, k^+^_Di_, k^-^_Di_ (5) T_i_ + LT ↔ TLT, k^+^_Ti_, k^-^_Ti_ (6) D_i_ + LT ↔ DLT, k^+^_Di_, k^-^_Di_ (7) T_i_ + LD ↔ TLD, k^+^_Ti_, k^-^_Ti_ (8) D_i_ + LD ↔ DLD, k^+^_Di_, k^-^_Di_ (9)TLD → DLT, k_cp_ (10) DLT → TLD, k_p_ (11) TLT → TLT’, k_t_ (12) TLT’ → TLT, k_t_ (13) DLD → DLD’, k_d_ (14) DLD → DLD, k_d_

Parameter values for the ANT model at Δɸ 180 mV: k^-^_Tm_ = 4×10^4^ s^−1^, k^+^ = 6.4×10^6^ M^−1^ s^−1^, k^-^_Ti_ = 200 s^−1^, k^+^ = 4×10^5^ M^−1^ s^−1^,k^-^_Dm_ = 4×10^4^ s^−1^, k^+^ = 4×10^6^ M^−1^ s^−1^, k^-^_Di_ = 100 s^−1^, k^+^ = 4×10^6^ M^−1^ s^−1^, k_p_ = 92 s^−1^, k_cp_ = 3.5 s^−1^, k_d_ = 4.8 s^−1^, k_t_ = 5.8 s^−1^

##### Modeled distribution of ANTs

The density of ANTs has been estimated at 0.2 nm/mg protein in rat liver mitochondria(Forman & Wilson, 1983). Assuming that 1 nm/mg protein is approximately 1.25 mM(Magnus & Keizer, 1997) leads to a concentration of 0.25 mM. With this concentration, the number of ANTs in a given reconstruction can be estimated. Using the total mitochondrial volume proportionality with ANT concentration, we set the number of ANTs in SFA2 as 7678. In the onion mitochondrion, the number of ANTs were set at 7531. Thus, both types of mitochondria analyzed contained a 17:1 ratio of ANTs to ATP synthases.

##### VDAC model

To model the exit of ATP molecules to the cytosol we included VDACs, the main mechanism for metabolites to cross the OM. We implemented a simple model assuming VDAC proteins interact with ATP molecules and translocate them to the cytosol by the reaction VDAC + ATP_IMS_ ⇌ VDAC + ATP_cyto_. In all simulations, VDAC proteins were homogeneously distributed in the OMM. VDAC abundances were set as proportional to the total mitochondrial volume encapsulated by the OMM.

Parameters for the VDAC mode: rate constant of the reaction, k_vdac_ = 1×10^6^ M^−1^ s^−1^, the density of VDACs(De Pinto *et al*, 1987), δ = 1×10^4^ μm^−2^, the number of VDACs considered in the simulations, N_vdac_ = 10268 for CM-containing SFA2 mitochondria and 4979 for onion-like SFA4 mitochondria.

##### Metabolite buffers

ATP and ADP molecules can interact with different cations, be bound, or ionized. The total concentration of ATP and ADP molecules can be distributed in several compounds like ATP^4−^, ADP^3−^, ATPMg^2−^, etc. The final distributions can be estimated by coefficients representing the fraction of unbound ATP in the matrix or the IMS. For our model, mitochondrial ADP^3−^ and ATP^4−^ concentrations were estimated analogously to published data (Magnus & Keizer, 1997) as [ADP]_m,free_ = 0.8 [ADP]_m_, [ATP]_m,free_ = [ATP]_m_, [ATP^4−^] = 0.05 [ATP]_free_ and [ADP^−3^] = 0.45 [ADP]_free_. The initial concentrations of ATP and ADP in the matrix were set to 13 mM and 2 mM, respectively, and to 6.5 mM and 0.1 mM in the IMS and cytosol. In some simulations, these concentrations were kept constant.

##### Well-mixed model of ATP generation

A system of ordinary differential equations was derived from the reactions above (given in(Garcia *et al*, 2022)) and used to calculate the rate of ATP generation in a well-mixed model, without considerations of mitochondrial geometry.

#### Molecular dynamics simulations

Coarse grained molecular dynamics models of systems with varying compositions were generated using insane.py (Wassenaar *et al*, 2015) and Martini 2.2 force-field parameters (Marrink *et al*, 2007, 2004). Minimization and equilibration followed the conventional protocols established by CHARMM-GUI (Qi *et al*, 2015; Jo *et al*, 2007), summarized briefly here. Initial soft-core minimization is followed by steepest descent to generate an integrator ready relaxed configuration. The systems are jumped to 303 K by random assignment of velocities and the systems undergo several steps of NPT restrained equilibration. All equilibration steps were run with the Berendsen barostat (Berendsen *et al*, 1984). Over the course of restrained equilibration stages, the timestep was gradually increased from 2 to 20 fs, and bilayer headgroup restraints were reduced from 200 to 20 KJ/mol nm^2^. Systems are followed by several microseconds of NPT production using a semiisotropic Parinello-Rahman barostat (Parrinello & Rahman, 1981). All equilibration and production simulations use the Bussi-Donadio-Parinello velocity-rescaling thermostat (Bussi *et al*, 2007) with reaction-field electrostatics and shifted Van der Waals potentials both with 1.1 nm cutoff (de Jong *et al*, 2016). Molecular dynamics simulations were run using gromacs 2022.1 (Bauer *et al*, 2022; Abraham *et al*, 2015). Force-field parameters, topologies, and simulation control parameters to reproduce this work are available https://github.com/RangamaniLabUCSD/2022-mitochondria-lipidomics-md. Henceforth all timescales reported are in simulation time, and not scaled using the conventional factor of 4 for Martini 2.2 simulations.

The bending modulus of the membrane for each composition was estimated by analyzing the height fluctuation spectra (Brown, 2008; Venable *et al*, 2015; Helfrich, 1973; Ergüder & Deserno, 2021; Fowler *et al*, 2016) of systems approximately 40 by 40 nm in size over 5 μs of production. Assuming the Helfrich Hamiltonian in the limit of small deformations (Monge gauge), zero membrane tension, and equipartition of energy, the power spectrum of the bilayer height fluctuations is given by,

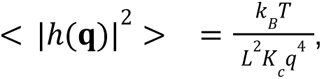

Where *q* is the magnitude of wave-vector **q**, *k_B_T* is the Boltzmann constant and temperature, *L* is the system length, and *K*_*c*_ is the membrane bending modulus. Quadrilateral meshes representing the surfaces of each leaflet are fit to the headgroup region (defined by the pointcloud of PO4 and GL0 beads) using piecewise cubic interpolation. The neutral surface of the bilayer was assumed to be the mean of the two surfaces. Computing the squared discrete fourier transform of the neutral surface, we obtained the 2D power spectrum of height fluctuations. The 2D power spectrum was converted into 1D for subsequent analysis by radially binning and averaging. Fitting the 1D power spectrum to the theoretical enables the estimation of the membrane bending modulus. Quantifying the error of the estimate is performed using parametric bootstrapping analysis following recommendations by Erguder and Deserno(Ergüder & Deserno, 2021). In brief, we have a sequence of mean squared amplitudes, <|*h*(**q**)|^2^> for each *q* corresponding to each trajectory frame. To obtain a meaningful error we consider these as samples from a continuous trajectory with some potential correlation. Statistical block averaging enables us to estimate the autocorrelation time of the data and further a correlation corrected standard deviation. The standard error of the bending modulus is determined from using parametric bootstrapping to sample values of spectral power for large wavenumbers. These are subject to a non-linear fit to obtain a distribution of *K* values for each system from which we obtain the standard deviation. Processing of the data for the analysis was performed using numpy (Harris *et al*, 2020), Scipy(Virtanen *et al*, 2020), MDAnalysis(Virtanen *et al*, 2020), and a modified curvature analysis framework(E., 2021) available from https://github.com/ctlee/membrane-curvature.

The local neighbor enrichment for each lipid type was investigated using MDAnalysis(Virtanen *et al*, 2020). For each lipid, we count the numbers and types of each lipid within a 1.5 nm radius. The position of each lipid was either the sole PO4 or GL0 bead in the headgroup region. Normalizing by the number of frames and copy number of each lipid type produces the mean number of lipids of each type around a lipid of a given type; Comparing this value against the probability derived from random chance with no interactions given by the system composition, we obtain the deviation from random chance.

Using smaller systems of approximately 15 by 15 nm in length we computed the lateral pressure profiles for each composition. Each small system was equilibrated for 4 μs followed by 200 ns of production with positions and velocities written out every 5 ps in full numerical precision. The stresses for each frame by reprocessing using gmx-ls in gromacs 2016.3(Vanegas *et al*, 2014). Contours for stress calculation were spaced approximately 1 nm in the X and Y directions (in-plane of the membrane) and 0.1 nm in the Z direction (normal to the membrane). The lateral pressure, *p*(*z*), is given by *p*(*z*) = σ*_ZZ_* − (σ*_XX_* + σ*_YY_*)/2, where σ*_XX_*, σ*_YY_*, and σ*_ZZ_* are the diagonal components of the stress tensor. The discrete lateral pressure profile was fit using a piecewise cubic interpolation and the bending moment, 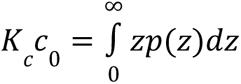, was evaluated using numerical integration of the interpolated function. Errors of values derived from and the lateral pressure profile were estimated by splitting the collected production frames into three non-overlapping chunks.

#### Continuum modeling of tubular cristae formation

##### Background

In the Helfrich-Canham-Evans model (Helfrich, 1973; Canham, 1970; Evans, 1973), membranes are treated as a two-dimensional surface with an elastic bending energy given by:

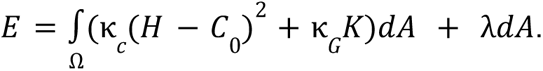

where *к_c_* is the bending modulus of the membrane (stiffness), *H* is the mean curvature of the structure, *C_0_* is the net spontaneous curvature across the bilayer, *к_G_* is the Gaussian modulus, *K* is the Gaussian curvature, and λ is the membrane tension. The total energy of the membrane (*E*) is obtained by integrating the energy density of the manifold over the area Ω. When this energy is minimized, the shape of the membrane corresponding to mechanical equilibrium is obtained.

##### Overview of the model

The mathematical derivations of this model can be found in extensive detail in (Mahapatra, 2022). Here, we provide a brief summary of the equations. The membrane is modeled as a thin elastic shell in mechanical equilibrium. Table 1 summarizes the symbols and notation; all nonscalar quantities are denoted by a bar overhead.

**Table 1:** Notation and list of symbols.

| Notation | Description | Unit |
| --- | --- | --- |
| $W$ | Free energy density of the membrane | pN/nm |
| $\kappa$ | Bending modulus of the membrane | pN · nm |
| $\kappa_d$ | Deviatoric modulus of the membrane | pN · nm |
| $\bar{n}$ | Surface normal | 1 |
| $a_{\alpha\beta}, a^{\alpha\beta}$ | Metric tensor and its contravariant | — |
| $b_{\alpha\beta}, b^{\alpha\beta}$ | Curvature tensor and its contravariant | — |
| $\bar{T}$ | Surface traction | pN/nm |
| $s$ | Arclength along the membrane | nm |
| $\psi$ | Angle made by surface tangent with the horizontal direction | 1 |
| $H$ | Mean curvature | nm<sup>-1</sup> |
| $D$ | Deviatoric curvature | nm<sup>-1</sup> |
| $C_0$ | Spontaneous mean curvature | nm<sup>-1</sup> |
| $D_0$ | Spontaneous deviatoric curvature | nm<sup>-1</sup> |
| $f^\alpha$ | Tangential component of the force per unit area | pN/nm<sup>2</sup> |
| $f_n$ | Normal component of the force per unit area | pN/nm<sup>2</sup> |
| $\lambda$ | Membrane tension | pN/nm |
| $\lambda_0$ | Tension at boundary | pN/nm |
| $F_n$ | Traction at the boundary | pN/nm |

The force balance on the membrane is given by

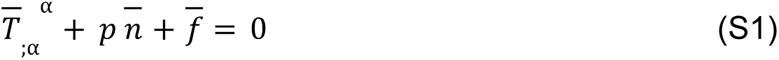

where, 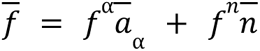 is the external force density applied to the membrane, *p* is normal pressure on the membrane and *T* is traction on the membrane and given by,

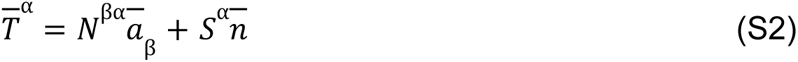

Here, *N̅* in-plane components of the stress and is given by

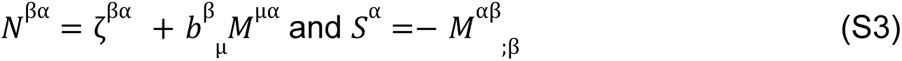

where, ζ^βα^ and ζ^βα^ are obtained from the following constitutive relationships (Steigmann, 2018)

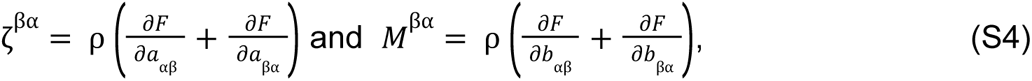

with *F* = *W*/ρ is the energy mass density of the membrane. Combining these we get the balance equations in tangent and normal direction

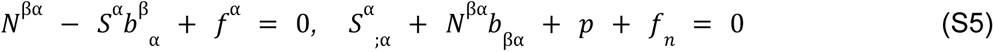

Here *f*^α^ and *f_n_* are the tangential and normal components of external force applied to the membrane per unit area.

The energy density of the membrane *W* is taken as follows to account for the mean and the deviatoric curvature

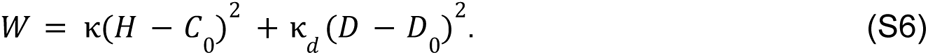

To obtain tubular shapes, recasting the Helfrich energy in terms of the isotropic spontaneous curvature *C_0_* and the anisotropic spontaneous curvature *D_0_*, is a commonly used approach(Kralj-Iglič *et al*, 2020; Kabaso *et al*, 2012; Mahapatra, 2022; Noguchi *et al*, 2022). In this case, the energy is written in terms of the mean curvature and the deviatoric curvature *D*. The deviatoric curvature is defined as half of the difference between the two principal curvatures.

The tangential force balance relation in Equation (S5)*_I_* simplifies as

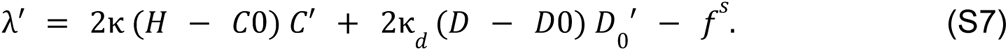

The normal force balance relation S5*_II_* (the shape equation) becomes

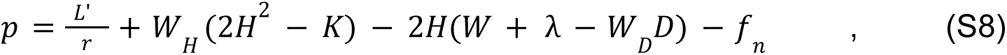

where *L* relates to the expression of the traction, given by

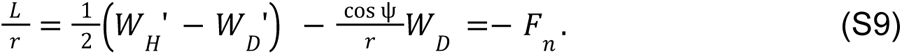

Where *F_n_* is the traction acting normal to the membrane. The above relation gives a natural boundary condition for *L* at both the boundaries. At the center, it directly correlates with the value of pulling force as

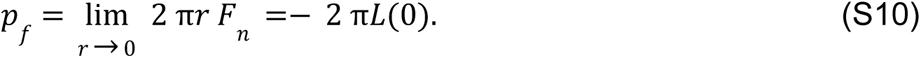

##### Area parameterization

The governing equations are solved in a patch of the membrane with fixed surface area, where the coat area of protein is prescribed. The arclength parametrization poses some difficulties since the total arclength varies depending on the equilibrium shape of the membrane. Therefore, we did a coordinate transformation of arclength to a local area *a* as given by

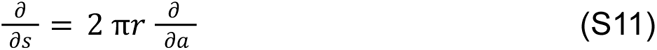

Note that in the differential form, local area relates as *da* = 2 π*r ds*

The tangential force balance relation in Equation S7 transforms to

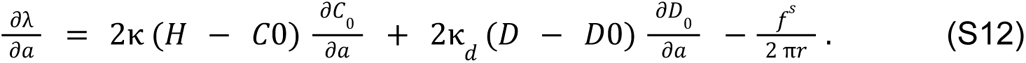

The normal force balance relation in Equation S8 becomes

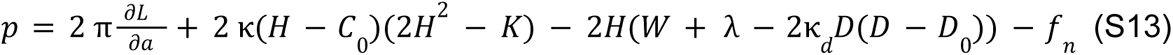

Where

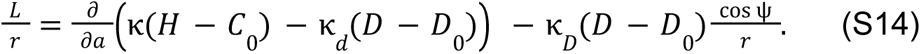

##### Numerical methods

We solved the system of equations (Equation S11 to Equation S14) numerically to get the equilibrium shape of the membrane for a coat of protein at the center of an axisymmetric area patch. The solution domain is presented in Figure S6A, along with the input protein coat and the boundary conditions shown in Figure S6A. The protein coat includes both the spontaneous mean curvature cap and a combination of mean and deviatoric spontaneous curvature in the rest of the coat region (Figure S6B). Note that we introduced a shape variable ψ, which denotes the angle made by the tangent from its radial plane. The membrane is clamped at the domain boundary, where both the displacement and the angle ψ = 0. The membrane tension is also prescribed at the boundary. At the pole, ψ is taken to be zero, which indicates the smoothness at the center of the membrane. *L* is set to zero, indicating that there is no pulling force acting at the center.

To solve the system of equations, we used MATLAB-based bvp4c, a finite difference-based ODE solver with fourth-order accuracy (MATLAB codes are available https://github.com/Rangamani-lab/arijit_deviatoric_tube.2022). We used a nonuniform grid ranging from 1000 to 10000 points, with the finer grid towards the center. We used a large domain size of 10^6^ nm^2^ to avoid boundary effects but we show the results focusing on the membrane deformation (region enclosed by the dashed line in Figure S6C). The values of the different parameters used are given in the table below.

List of parameters used in the simulations.

| Notation | Description | Range |
| --- | --- | --- |
| $C_0$ | Spontaneous mean curvature | 0.01–0.035 $\text{nm}^{-1}$ |
| $D_0$ | Spontaneous deviatoric curvature | 0–0.035 $\text{nm}^{-1}$ |
| $\kappa = (\kappa_d)$ | Bending modulus of the membrane | 10–20 $k_B T$ |
| $a_{coat}$ | Coat area of protein | $1.413 \times 10^4 \text{ nm}^2$ |
| $a_{mem}$ | Total area of the membrane | $5.65 \times 10^5 \text{ nm}^2$ |
| $\lambda_0$ | Tension at boundary | 0.01 pN/nm |

##### Parameter ranges

The parameter ranges for the continuum model were chosen from the literature and from the CGMD simulations. The range of anisotropic spontaneous curvature induced by the ATP synthases was estimated from the CGMD simulations presented in (Anselmi *et al*, 2018) for a single ATP synthase dimer by estimating the two principal curvatures for the small deformation seen in Figure 1 of that work. Note that these estimates are obtained from digitizing the images and do not contain the information carried in the thermal fluctuations. The bending moduli range was in the range consistent with CG-MD calculations and previous experimental measurements. The tension and the coat area are free parameters in the model and were tuned such that we could obtain tubules of length and radius consistent with experimental measurements as shown in Figure 5B.

